# The human non-visual opsin OPN3 regulates pigmentation of epidermal melanocytes through interaction with MC1R

**DOI:** 10.1101/552851

**Authors:** Rana N. Ozdeslik, Lauren E. Olinski, Melissa M. Trieu, Daniel D. Oprian, Elena Oancea

## Abstract

Opsins form a family of light-activated, retinal-dependent G-protein coupled receptors (GPCRs) that serve a multitude of visual and non-visual functions. Opsin3 (OPN3 or encephalopsin), initially identified in the brain, remains one of the few members of the mammalian opsin family with unknown function and ambiguous light-absorption properties. We recently discovered that OPN3 is highly expressed in human epidermal melanocytes—the skin cells that produce melanin. The melanin pigment is a critical defense against ultraviolet radiation and its production is mediated by the Gαs-coupled melanocortin-1 receptor (MC1R). The physiological function and light-sensitivity of OPN3 in melanocytes is yet to be determined. Here we show that in human epidermal melanocytes OPN3 acts as a negative regulator of melanin production by interacting with MC1R and modulating its cAMP signaling. OPN3 negatively regulates the cAMP response evoked by MC1R via activation of the Gαi subunit of G-proteins, thus decreasing cellular melanin levels. In addition to their functional relationship, OPN3 and MC1R colocalize at both the plasma membrane and in intracellular structures and form a physical complex. Remarkably, OPN3 can bind retinal, but does not mediate light-induced signaling in melanocytes. Our results identify a novel function for OPN3 in the regulation of the melanogenic pathway in epidermal melanocytes. Our results reveal a light-independent function for the poorly characterized OPN3 and a novel pathway that greatly expands our understanding of melanocyte and skin physiology.

**Significance:** Our data reveals a novel function for the non-visual opsin OPN3 in regulating the pigmentation of human melanocytes by interacting with and modulating the activity of MC1R.

## Introduction

Unlike most mammals which have melanin-producing melanocytes predominantly in the hair follicle bulb [1], humans are uniquely equipped with melanocytes in the outermost layer of the skin, the epidermis [2, 3]. These neural crest-derived melanocytes are the only source of the photoprotective pigment melanin in human skin and, thus, are critical for the defense against solar ultraviolet radiation (UVR)-induced genotoxic damage [4–6].

Solar UVR at the surface of the earth is comprised of ~5% short-wavelength UVB and ~95% long-wavelength UVA. Much of our current knowledge about melanogenesis in epidermal melanocytes stems from the well-characterized UVB-induced melanin pathway [7]. UVB elicits DNA damage in epidermal keratinocytes, triggering facultative skin darkening through increased melanin production in neighboring melanocytes [8]. UVB irradiated keratinocytes and melanocytes locally secrete alpha-melanocyte stimulating hormone (αMSH), an agonist of the Gαs-coupled melanocortin-1 receptor (MC1R) that is primarily expressed on melanocytes [8, 9]. MC1R has a pivotal role in determining pigmentation, as several naturally occurring loss-of-function MC1R variants are associated with the “red hair phenotype” [10, 11] characterized by pale complexion and increased sensitivity to UVR [12]. Downstream, αMSH-induced MC1R activation leads to stimulation of adenylyl cyclase (AC) and production of cyclic adenosine monophosphate (cAMP). Accumulation of cAMP, through several molecular steps, induces upregulation of microphthalmia transcription factor (MITF)—the master transcription factor leading to increased expression of melanogenic enzymes like tyrosinase (TYR) [13].

In addition to the UVB pathway, we recently characterized a novel UVA-induced melanogenic pathway in human epidermal melanocytes (HEMs). This retinal-dependent phototransduction pathway is mediated by Gαq/11 activation, resulting in a rapid increase in intracellular Ca^2+^ and elevated cellular melanin levels [14–16]. The identity of the putative GPCR that mediates the UVA phototransduction cascade remains unknown, but members of the retinal-dependent, light-sensitive opsin family are ideal candidates. Coincidentally, in HEMs, we and others have found expression of mRNA corresponding to several opsins [16–20]; opsin3 (OPN3), which has unknown physiological function, has significantly higher mRNA expression than any other detected opsin [18].

The scarcity of functional data on OPN3, discovered nearly 20 years ago [21, 22], may derive from its unique structure and widespread expression ranging from deep brain regions [21, 22] to peripheral tissues [23]. OPN3 has a unique, long carboxy (C)-terminus with no sequence homology to any known GPCR. Recent studies have determined the photoreceptive properties for several OPN3 homologs [24–26]: zebrafish, pufferfish, and chicken OPN3 absorb blue light (λ_max_=465 nm) [25]; mosquito OPN3 forms a bistable photopigment with 13-*cis*, 11-*cis* and 9-*cis* retinal, absorbs blue-green light (λ_max_=490 nm), and activates G-proteins Gαi/o in a light-dependent manner [24, 26]. *However, the light sensitivity, G-protein coupling, and function of human OPN3 remain unknown*.

Here we show that OPN3 is a negative regulator of melanogenesis in human melanocytes. OPN3 does not mediate the UVA-evoked Ca^2+^-response of HEMs, nor does it modulate the sensitivity of these cells to visible light, despite being able to bind 11-*cis* and all-*trans* retinal. OPN3 couples to Gαi to negatively regulate the αMSH-induced cAMP response of MC1R. In addition, OPN3 and MC1R form a physical complex. Our data identify a novel melanogenic regulatory mechanism and a key function of human OPN3 in skin, both of which expand our knowledge of melanocyte physiology.

## Results

### OPN3 does not mediate Ca^2+^-dependent UVR phototransduction in HEMs

Physiological doses of UVR induce a retinal- and PLCβ-dependent transient increase in cytosolic Ca^2+^, mediated by activation of Gαq/11 via an unknown putative GPCR [14–16]. Because mosquito OPN3 activates Gαi/o subunits of G-proteins in a light-dependent manner [24] and the Gβγ subunits that dissociate from Gαi are able to activate PLCβ and cause a Ca^+2^ response, we reasoned that OPN3 may be the GPCR that mediates UVR phototransduction in HEMs. Like all opsins, OPN3 possesses a lysine in the seventh transmembrane domain (K299) and a negatively-charged counterion in the third transmembrane domain (D117) (**Fig. 1A**), both involved in the binding of retinal chromophore to the opsin apoprotein [27], thus conferring light sensitivity.

**Figure 1.**
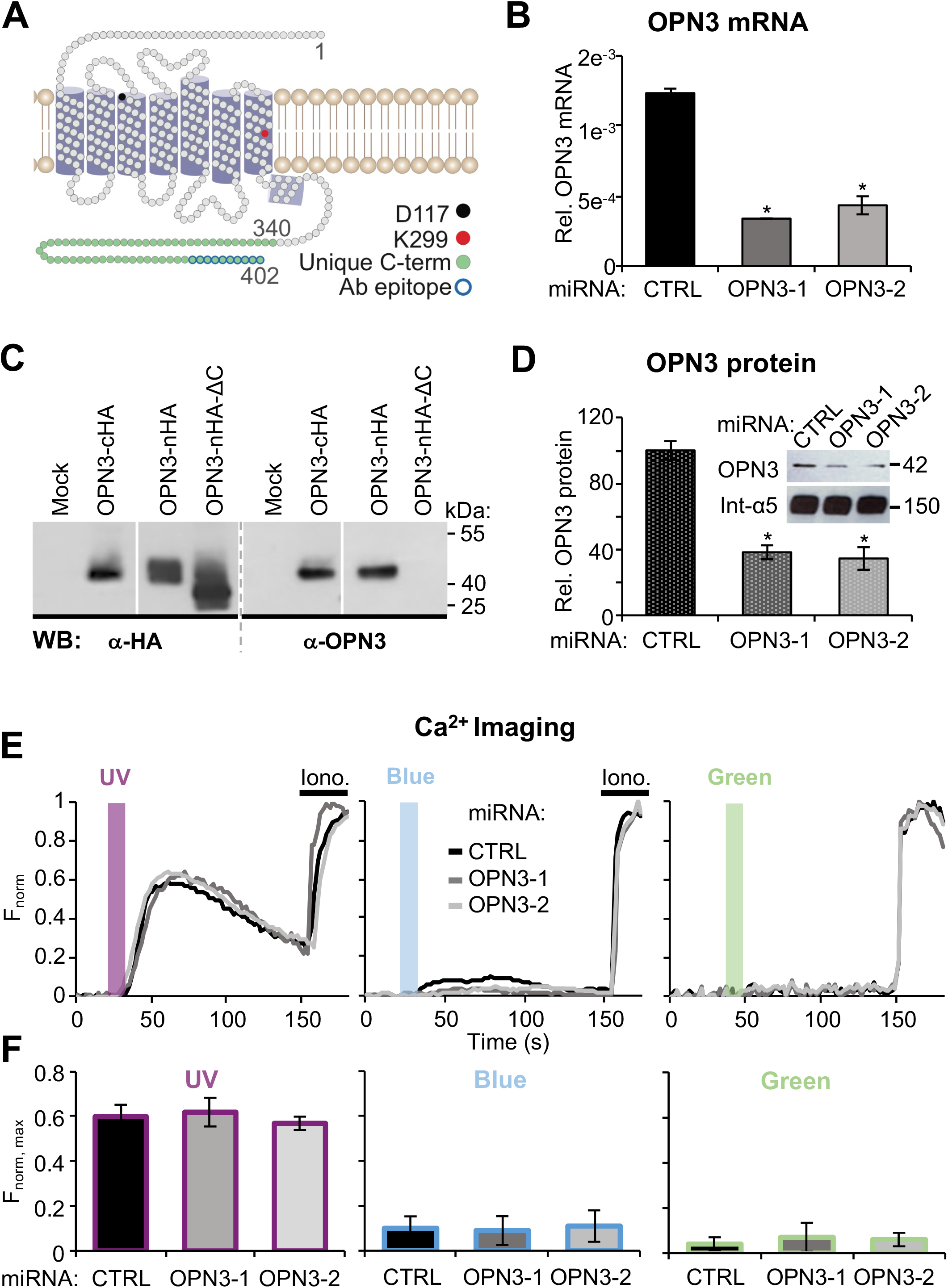
Human OPN3 does not mediate light induced Ca^2+^ responses in HEMs. **A. Two-dimensional model of human OPN3 (hOPN3).** hOPN3, similar to rhodopsin, has 7 predicted transmembrane domains followed by a short intracellular helix (purple cylinders) and a unique C-terminus (amino acids 340-402, green) containing the epitope for the anti-OPN3 antibodies (blue outline) [58]. Each circle represents one amino acid. The conserved lysine K299 (red) is involved in retinal binding, which forms a Schiff base with counterion D117 (black). **B. OPN3 mRNA levels in HEMs expressing control (CTRL) or OPN3-targeting miRNAs (OPN3-1 or OPN3-2).** QPCR analysis of OPN3 relative to actin mRNA levels. OPN3-1 and OPN3-2 miRNA expressing HEMs have decreased OPN3 mRNA levels by ~70% and ~60% respectively, compared to control miRNA transduced HEMs. n=3 independent experiments, ± SEM. * p< 0.01 **C. Specificity of the anti-OPN3 antibody.** OPN3 tagged with HA at the C-terminus (OPN3-cHA) or N-terminus (OPN3-nHA), N-terminal HA-tagged OPN3 mutant missing the last ten amino acids (OPN3-nHA-ΔC), or empty vector (Mock) were expressed in HEK293 cells. Immunoblots using anti-HA or anti-OPN3 antibodies show the same size band (~42 kDa) corresponding to OPN3 (calculated molecular weight 45 kDa) for both full-length constructs but not for the ΔC mutant, which resulted in a ~40 kDa band using anti-HA and no band using anti-OPN3 antibodies. Representative of n=4 independent experiments. **D. OPN3 protein levels in HEMs expressing control (CTRL) or OPN3-targeting miRNAs (OPN3-1 or OPN3-2).** Densitometric analysis of HEMs expressing CTRL, OPN3-1 or OPN3-2 miRNA and immunoblotted with anti-OPN3 or anti-integrin α5 antibodies (inset). Bars represent OPN3 protein level normalized to integrin α5. n=3 independent experiments, ± SEM. * p< 0.01. **E and F. Light-induced Ca^2+^ signaling in HEMs is not dependent on OPN3 expression. E.** Fluorescent Ca^2+^ imaging of HEMs expressing CTRL, OPN3-1 or OPN3-2 miRNA and stimulated with 200 mJ/cm^2^ ultraviolet (UV, λ_max_=360 nm), blue (λ_max_=450 nm) or green (λ_max_=550 nm) light, normalized to the maximal Fluo4-AM Ca^2+^ response obtained with ionomycin (Iono). Each trace represents the average of 10-20 cells from one coverslip. **F.** Average amplitude of Ca^2+^ responses of HEMs under the conditions shown in **E.** n=5 independent experiments for each bar, ± SEM.

To determine if OPN3 mediates UVR phototransduction, we reduced *OPN3* levels in HEMs using two OPN3-targetted miRNAs (OPN3-1 and OPN3-2). Each miRNA reduced the level of *OPN3* mRNA by more than 60% compared to control scrambled miRNA (CTRL) (**Fig. 1B**). To confirm that OPN3 protein levels were also reduced, we first validated an antibody against the unique C-terminus of OPN3 (**Fig. 1C**). Using this antibody, we showed that OPN3 protein levels are also reduced by more than 60% in HEMs expressing OPN3-1 or OPN3-2 miRNA (**Fig. 1D**). We monitored intracellular Ca^2+^ levels using the fluorometric Ca^2+^ indicator Fluo-4 AM in HEMs preincubated with all-*trans* retinal and expressing OPN3-1, OPN3-2 or CTRL miRNAs. Exposure to UVR (200 mJ/cm^2^) led to a synchronized and transient Ca^2+^ response of similar amplitude in both HEMs expressing CTRL or OPN3-1 or OPN3-2 miRNAs (**Figs. 1E and 1F**, left panels), indicating that OPN3 is not required for the UVR-induced Ca^2+^ response.

We next determined if OPN3 mediates blue or green light-induced Ca^2+^ changes in HEMs since it does not regulate the UVR-Ca^2+^ cascade. Although we have previously shown in HEMs that the UVR-Ca^2+^ photocascade is UVR specific and no Ca^2+^ responses could be evoked by similar doses of visible light [16], we questioned if human OPN3 could act to suppress the Ca^2+^ response to blue-green light in HEMs. In this case, reducing OPN3 expression may uncover a measurable Ca^2+^ response to blue or green light. This idea is plausible given that OPN3 homologues are sensitive to blue-green light [24]. We monitored the Ca^2+^ responses of HEMs expressing CTRL or OPN3-targeted miRNAs, pre-incubated with all-*trans* retinal, and exposed to 200 mJ/cm^2^ of blue (λ_max_=450 nm) or green (λ_max_=550 nm) light. HEMs expressing CTRL miRNA did not have a significant Ca^2+^ response to blue or green light, and neither did HEMs expressing OPN3-targetted miRNAs (**Figs. 1E and 1F**, middle and right panels).

Because the miRNAs partially reduced OPN3 expression and the residual OPN3 could be sufficient for the UVR response, we used CRISPR/Cas9 to eliminate OPN3. Primary HEMs with CRISPR/Cas9-induced mutations did not survive the selection of clonal lines; we used immortalized human epidermal melanocytes (Hermes 2b), which express similar levels of MC1R and OPN3 mRNA as HEMs (**Supp. Fig. 1A**) to generate clonal CRISPR/Cas9 lines. Hermes 2b cells containing a mutation introduced in exon1 of OPN3 lack OPN3 expression (**Supp. Fig. 1B**). We performed Ca^2+^ imaging experiments in wild-type Hermes 2b cells or those lacking OPN3 expression and obtained similar results to Figs. 1E and 1F: the UVR-induced Ca^2+^ response was not affected by the absence of OPN3 expression and no responses to blue or green light were measured with or without OPN3 (**Supp. Figs. 1D, E**). These results further bolster our conclusion that OPN3 does not mediate Ca^2+^-dependent phototransduction of UVR, blue, or green light in melanocytes.

These findings raised an interesting question: does human OPN3 form a photopigment? Spectroscopic analyses of recombinant OPN3 homologues indicated that they can bind 11-*cis* retinal and have an absorption maximum at ~470 nm [24, 25]. To determine if human OPN3 and retinal form a photopigment, we expressed C-terminal truncated, 1D4-tagged human OPN3 (OPN3ΔC-c1D4) [28] in HEK293 GnTI^−^ cells. We also expressed a variant in which the retinal-binding residue K299 was mutated to glycine [OPN3(K299G)ΔC-c1D4] (**Fig. 2A**, right), a mutation shown to abolish retinal-binding and light sensitivity of rhodopsin and other opsins [29, 30]. Neither the OPN3 C-terminal truncation nor the K299G mutation affected the cellular localization of OPN3 in a heterologous system as compared to C-terminal-tagged, full-length OPN3-MCherry (OPN3-cMCh) (**Fig. 2B; Fig. 2A**, left). UV-visible spectroscopy using purified OPN3ΔC-c1D4 protein or the K299G mutant protein revealed a significant protein peak for both proteins, but no detectable absorption peak at λ_max_>300 nm for either (**Fig. 2C**). To test if retinal was bound to human OPN3, we treated the purified proteins with a mixture of hydroxylamine (NH_2_OH) and sodium dodecyl sulfate (SDS) (**Fig. 2C**, insets). When retinal is bound to the apoprotein, SDS denatures the protein to expose the Schiff base that reacts with NH_2_OH to release retinal oxime, which absorbs at λ_max_=360 nm [31, 32]. Upon treatment with NH_2_OH and SDS, a retinal oxime absorption peak was observed for OPN3ΔC-c1D4 treated with both 11-*cis* retinal (Fig. 2C, left) or all-*trans* retinal (**Supp. Fig. 2A**) but not for OPN3(K299G)ΔC-c1D4 (**Fig. 2C**, right). This data suggests that OPN3, unlike the K299G mutant, can bind 11-*cis* retinal and all-*trans* retinal. Moreover, the reduced amplitude of the retinal oxime peak compared to the protein peak (λ_max_=280 nm) and purity of protein samples (**Supp. Fig. 2C**) suggest that OPN3 binds retinal in a *weak* manner.

**Figure 2.**
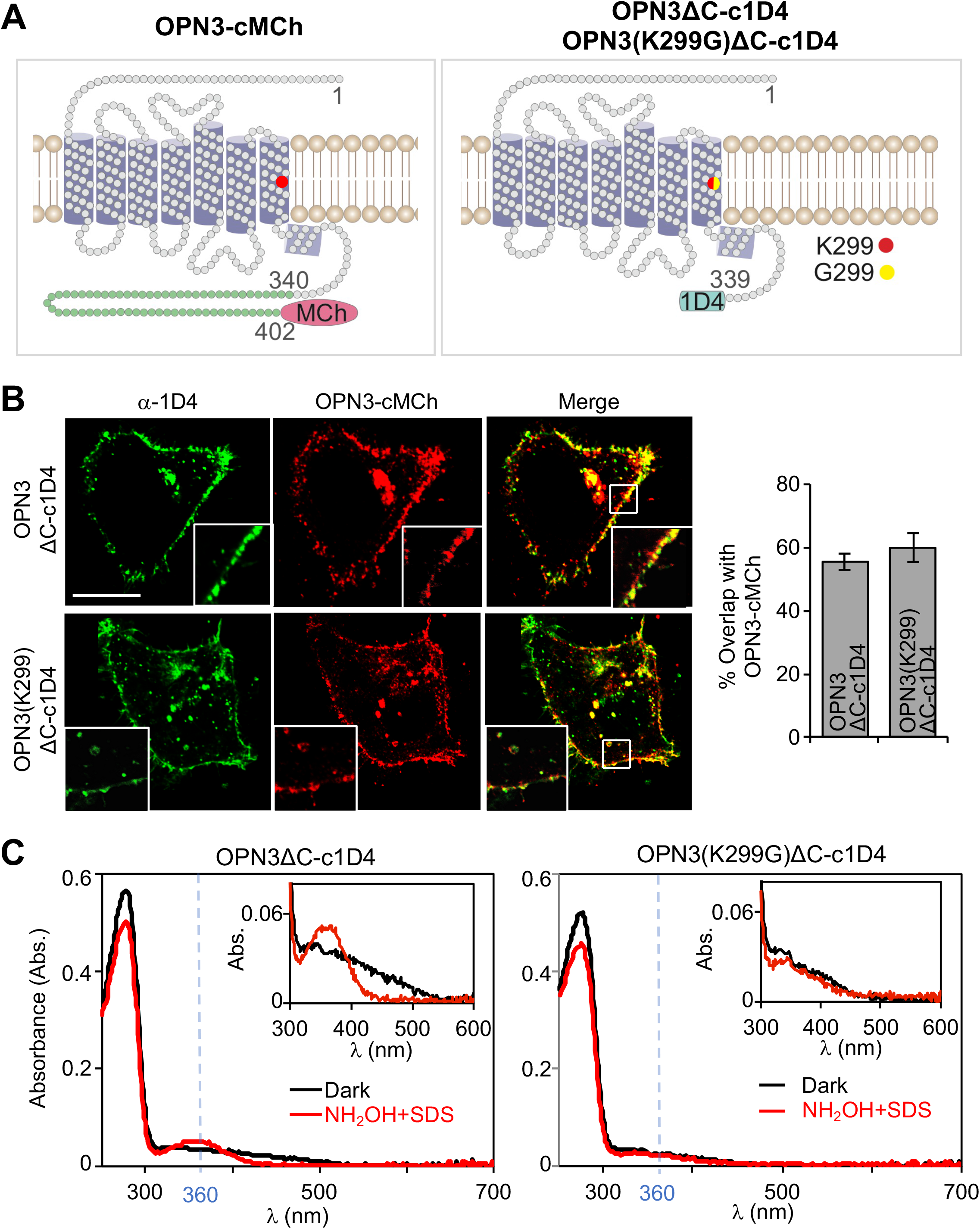
OPN3 requires the K299 residue for retinal binding. **A. Schematic representation of OPN3 variants.** We purified 1D4-tagged OPN3 with partially truncated C-terminus (OPN3ΔC-c1D4) or a K299G mutation of this variant known to inhibit retinal binding (OPN3(K299G)ΔC-c1D4). **B. OPN3ΔC-c1 D4 and OPN3(K299G)ΔC-c1D4 maintain the cellular localization of full-length OPN3.** Confocal images of HeLa cells co-expressing OPN3-cMCh and OPN3ΔC-c1D4 or OPN3(K299G)ΔC-c1D4 and immunostained with anti-1D4 antibody. OPN3ΔC variants have similar cellular localization as OPN3-cMCh. Quantitative analysis of OPN3-cMCh colocalization with OPN3ΔC-c1D4 or OPN3(K299)ΔC-c1D4, measured as percent overlap between the two fluorescent signals, shows significant colocalization (bar graph). Calibration bar: 10 μm; n=30 cells from 3 independent experiments. **C. UV-visible absorption spectra of OPN3ΔC and OPN3(K299G)ΔC.** Absorption spectra of purified OPN3ΔC-c1D4 and OPN3(K299G)ΔC-c1D4 incubated in 11-*cis* retinal were measured in the dark (black) and after hydroxylamine (NH**2**OH) and sodium dodecyl sulfate (SDS) treatment (red). Absorption spectra measured in the dark have similar protein peaks at λ_max_=280 nm for the two OPN3 variants. NH_2_OH+SDS treatment of OPN3ΔC-c1D4, but not OPN3(K299G)ΔC-c1D4, led to a peak at λ_max_=360 nm corresponding to retinal oxime. Inset: The retinal oxime peak of OPN3ΔC-c1D4 was approximately 10 times smaller than protein peak. Traces representative of n=3 independent experiments.

### OPN3 is a negative regulator of melanin levels in human melanocytes

We observed that HEMs with reduced OPN3 expression (OPN3-1 and OPN3-2 miRNAs) appeared more pigmented than CTRL miRNA-expressing HEMs (**Fig 3A**, left insets). Quantification of cellular melanin revealed that HEMs with reduced OPN3 had significantly higher melanin levels than control (**Fig 3A**, left graph). Similar to miRNA-treated HEMs, Hermes 2b cells lacking OPN3 have significantly higher melanin levels than control cells (**Supp. Fig. 1C**). We then tested the reciprocal: do elevated OPN3 levels lead to reduced melanin? Because we were not able to efficiently express exogenous OPN3 in HEMs, we used the pigmented melanoma cell line MNT-1. MNT-1 cells are highly pigmented and have been used extensively as a model system for melanocyte function because they preserve most primary melanocyte signaling and trafficking pathways. MNT-1 cells have MC1R levels comparable to HEMs, but lower OPN3 mRNA levels (**Fig. 3B**). We transfected MNT-1 cells with OPN3-cMCh or with only MCherry (MCh) and quantified cellular melanin levels. MNT-1 cells expressing OPN3-cMCh have an approximate 60% reduction in melanin levels as compared to MNT-1 expressing MCh (**Fig. 3A**, right graph). These results indicate that OPN3 negatively modulates the melanin levels of human melanocytes.

**Figure 3.**
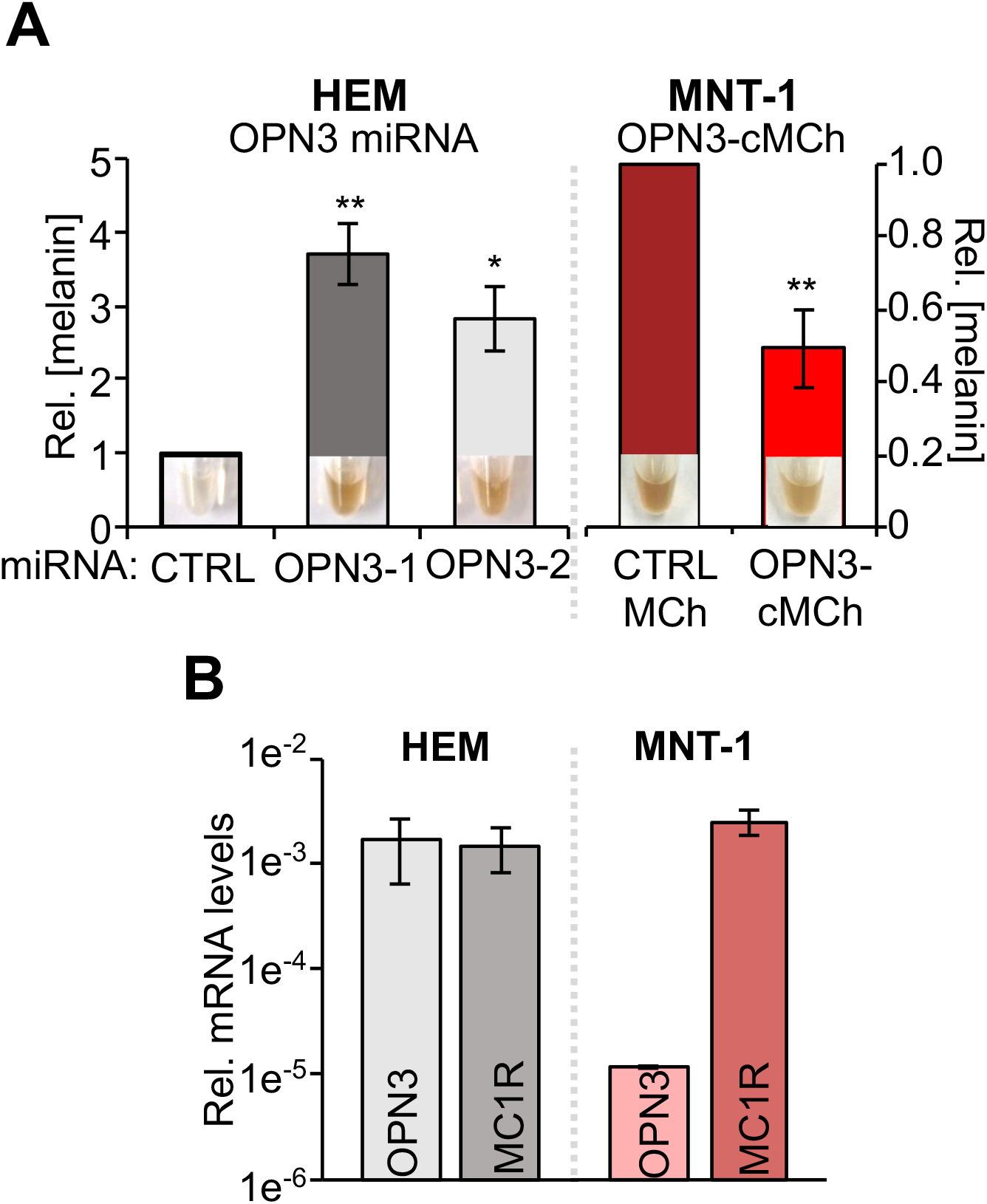
OPN3 expression inversely correlates with cellular melanin concentration. **A. Effect of OPN3 levels on melanin.** HEMs expressing OPN3-1 or OPN3-2 miRNA have significantly higher melanin levels compared to CTRL expressing cells (left), while MNT-1 cells expressing OPN3-MCh have reduced melanin compared to control MCh-expressing cells (right). Insets: representative pellets from each condition reflecting melanin levels. n=3 independent experiments, ± SEM. **B. OPN3 and MC1R mRNA levels in HEM and MNT-1.** mRNA levels of OPN3 and MC1R in HEMs and MNT-1 cells were measured by qPCR relative to actin. MNT-1 cells have similar MC1R levels as HEMs, but significantly lower OPN3 expression. n=3 independent experiments, ± SEM. * p< 0.05, ** p< 0.01.

### OPN3 is a negative regulator of MC1R-mediated signaling in human melanocytes

Basal melanin levels are regulated by MC1R; αMSH stimulates Gαs-coupled MC1R to increase cAMP levels. Ultimately, this cascade upregulates microphtalmia-associated transcription factor (MITF) which increases levels of the main melanogenic enzyme tyrosinase (TYR), and results in increased cellular melanin. Because mosquito OPN3 couples to Gαi/o G-proteins and Gαi/o generally signals by decreasing cellular cAMP, we tested if the negative effect of OPN3 on pigmentation is due to its inhibition of MC1R-mediated cAMP signaling.

To measure changes in cellular cAMP levels we used the validated FRET-based, genetic cAMP sensor Epac H187 [33] (**Supp. Fig. 3**). MNT-1 cells transfected with Epac H187 and either OPN3-cMCh or MCh were stimulated with αMSH. MNT-1 expressing MCh had a 50% increase in cellular cAMP measured as the change in FRET ratio (CFP/YFP), normalized to the maximum cAMP signal elicited by a mixture of adenylyl cyclase activator forskolin (FSK) and phosphodiesterase inhibitor 3-isobutyl-1-methylxanthine (IBMX) (**Figs. 4Ai** red trace and **4Aiv**). In contrast, MNT-1 cells expressing exogenous OPN3-MCh stimulated with αMSH had a significantly smaller increase in cellular cAMP levels (**Figs. 4Ai** dark blue trace and **4Aiv**), suggesting that OPN3 attenuates MC1R-mediated cAMP signaling in MNT-1 cells. We performed the same cAMP experiments in Hermes 2b cells expressing C-terminal YFP-tagged OPN3 (OPN3-cYFP) or YFP alone along with the red fluorescence-based cAMP indicator, R-FlincA [34]. Because Hermes 2b cells have lower levels of MC1R (**Supp. Fig. 1A**), we used the more sensitive cAMP indicator R-FlincA instead of Epac H187. Hermes 2b cells expressing YFP (CTRL YFP) exhibit a significant αMSH-induced cAMP response, while expression of OPN3-cYFP leads to a negligible cAMP response under similar conditions. These results confirm that OPN3 negatively modulates αMSH–induced MC1R cAMP signaling (**Fig. 4Ci** yellow v. light green traces **and 4Cii**).

**Figure 4.**
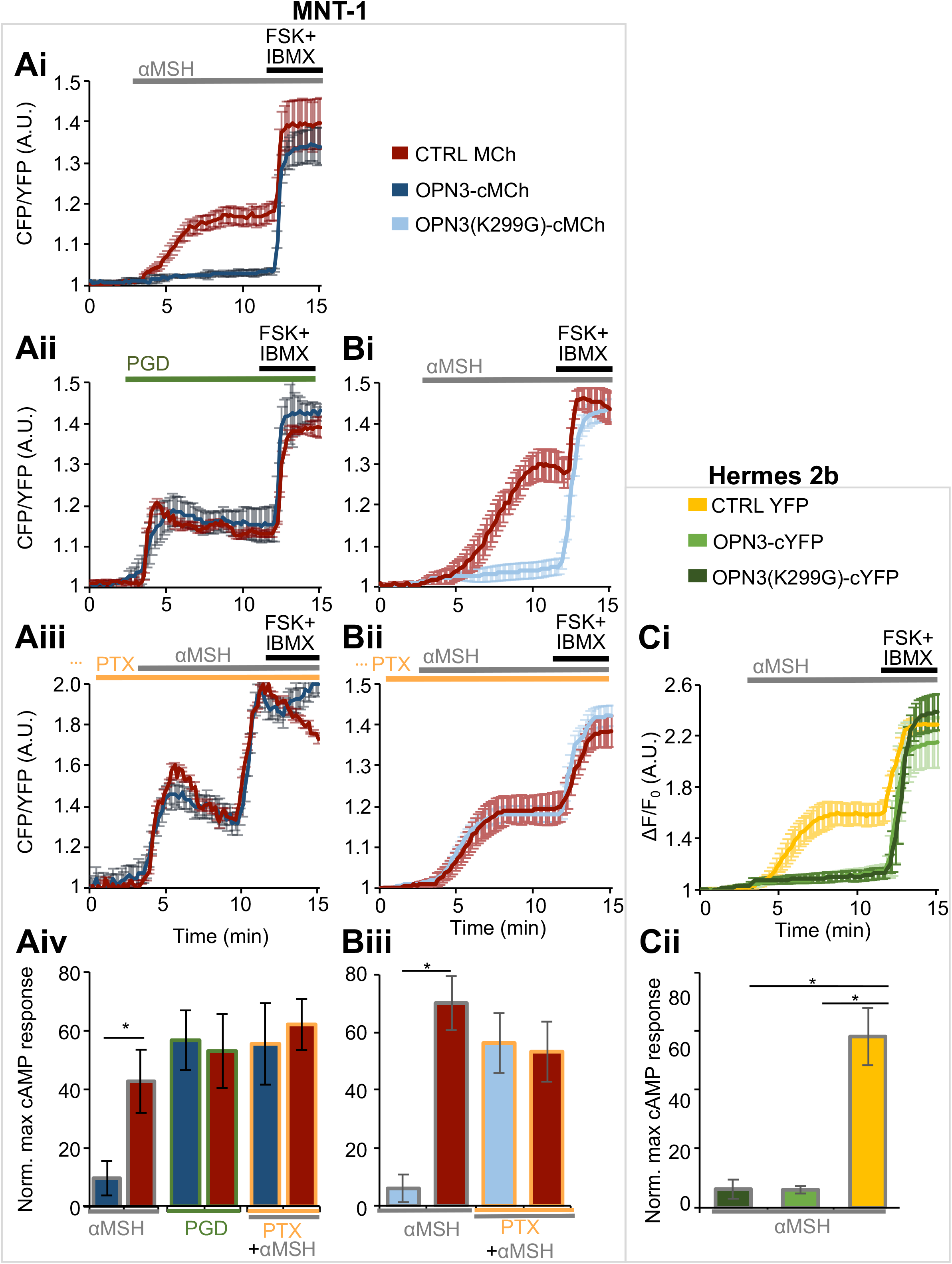
OPN3 modulates MC1R signaling. **A. OPN3 inhibits the MC1R-evoked cAMP response of via a PTX-sensitive mechanism in MNT-1 cells.** **Ai. OPN3 inhibits the αMSH-induced cAMP responses of MC1R.** MNT-1 cells expressing the FRET-based cAMP indicator Epac H187 and OPN3-cMCh or MCh alone (CTRL) were stimulated with αMSH (1 μM). The cAMP response of individual cells was monitored as the ratio of CFP and YFP fluorescence intensities and represented as a function of time. FSK and IBMX were added to elicit a maximal cAMP response, used for normalization. αMSH elicits a significant cAMP response in cells expressing MCh (CTRL, red trace), but not in cells expressing OPN3-cMCh (blue trace). n=5-10 cells, ± SEM. **Aii. OPN3 specifically attenuates the MC1R-mediated cAMP response.** Prostaglandin (PGD, 5 μM)-mediated activation of the endogenous prostaglandin E2 receptor leads to an increase in cellular cAMP (red trace) that is not attenuated in the presence of OPN3-cMCh (blue trace). n=5-10 cells, ± SEM. **Aiii. OPN3-mediated attenuation of MC1R signaling is pertussis toxin (PTX) sensitive.** MNT-1 cells treated with PTX (200 ng/ml, 4 h), which specifically inhibits the Gαi subunit of G proteins, and stimulated with αMSH exhibited a similar cAMP response in both MCh (red trace) and OPN3-cMCh (blue trace) expressing cells. n=5-10 cells, ± SEM. **Aiv. OPN3 inhibits MC1R signaling specifically and in a Gαi-dependent manner.** Average normalized amplitudes of αMSH- or PGD-induced cAMP responses in the presence or absence of PTX show that OPN3 expression significantly reduces the amplitude of cAMP responses to αMSH, but not PGD, and this effect is prevented by preventing Gαi activation with PTX. n=3 independent experiments per condition, ± SEM. * p< 0.05 **B. OPN3 inhibits MC1R-evoked cAMP responses independent of its retinal-binding ability.** **Bi. The OPN3-mediated inhibition of MC1R-induced cAMP responses does not require retinal-binding.** MNT-1 cells expressing OPN3(K299G)-cMCh (blue trace) and stimulated with αMSH (1 μM) exhibited significantly lower cAMP responses compared to MCh-expressing cells (red trace). n=5-10 cells, ± SEM. **Bii. OPN3(K299G)-mediated attenuation of MC1R signaling is PTX sensitive.** MNT-1 cells treated with the Gαi inhibitor PTX (200 ng/ml, 4 h) and stimulated with αMSH exhibited a similar cAMP response in both MCh (red trace) and OPN3(K299G)-cMCh (blue trace) expressing cells. n=5-10 cells, ± SEM. **Biii. OPN3(K299G) inhibits MC1R signaling in a Gαi-dependent manner.** Average normalized amplitudes of αMSH-induced cAMP responses of OPN3(K299G)-cMCh or MCh expressing MNT-1 cells. OPN3(K299G)-cMCh expression significantly reduces the amplitude of cAMP responses. n=3 independent experiments per condition, ± SEM. * p< 0.01 **C. OPN3 inhibits MC1R-evoked cAMP responses in Hermes 2b immortalized human epidermal melanocytes independent of retinal-binding.** **Ci. OPN3 inhibits the αMSH-induced cAMP responses of MC1R in Hermes 2b melanocytes in a retinal-independent manner.** Hermes 2b cells expressing the red fluorescent cAMP indicator R-FlincA and OPN3-cYFP, OPN3(K299G)-cYFP or YFP alone (CTRL) were stimulated with αMSH (1 μM). αMSH elicits a significant cAMP response in cells expressing YFP (CTRL, yellow trace), but not in cells expressing OPN3(K299G)-cYFP (light green trace) and OPN3-cYFP (dark green trace). n=5-10 cells per condition, ± SEM. **Cii. OPN3 and OPN3(K299G) inhibit MC1R signaling in Hermes 2b cells.** Average normalized amplitudes of αMSH-induced cAMP responses of Hermes 2b expressing OPN3-cYFP, OPN3(K299G)-cYFP are significantly lower than for YFP alone (CTRL). n=2 independent experiments per condition, ± SEM. **p < 0.01.

To determine whether OPN3 modulation is specific to MC1R-mediated cAMP signaling, we stimulated the endogenous Gαs-coupled prostaglandin EP2 receptor in MNT-1 cells expressing MCh or OPN3-cMCh. Stimulation of the EP2 receptor with prostaglandin led to a ~60% increase in cAMP both in MNT-1 cells expressing OPN3-cMCh and those expressing MCh (**Figs. 4Aii and 4Aiv**). This suggests that OPN3 does not modulate all Gαs-coupled receptors, but specifically regulates MC1R-mediated cAMP signaling.

We next determined how OPN3 negatively regulates MC1R signaling. Gαs signaling can be inhibited by the act of a Gαi-coupled GPCR [35]. To test if OPN3 couples to Gαi to reduce MC1R-mediated cAMP accumulation, we treated MNT-1 cells expressing MCh or OPN3-cMCh with Pertussis toxin (PTX), an inhibitor of Gαi signaling [8]. PTX-treated MNT-1 cells expressing MCh had a robust cAMP response to αMSH, as expected. PTX-treated MNT-1 expressing OPN3-cMCh had a similarly robust response to αMSH, unlike the negative regulatory affect OPN3 displays without PTX (**Figs. 4Aiii** compared to 4Ai **and 4Aiv**). PTX-mediated inhibition of Gαi signaling abolishes the negative regulation OPN3 has on MC1R cAMP signaling and suggests that OPN3 couples to Gαi. Interestingly, the lysine mutation of OPN3(K299G) did not affect the OPN3-mediated negative modulation of MC1R signaling (**Figs. 4Bi and 4Biii**) and PTX abolished the effect of OPN3(K299G) in MNT-1 and Hermes 2b (**Figs. 4Bii, 4Biii, 4Ci, 4Cii**). This indicates that OPN3 has a regulatory influence over MC1R despite its ability to bind retinal.

After MC1R-mediated cAMP levels increase, several molecular steps, including increased expression of MITF and upregulation of TYR, lead to an increase in melanin levels [36]. If OPN3 negatively regulates this pathway, as our cAMP data suggests, reducing OPN3 expression should increase MC1R-mediated signaling, increasing MITF and TYR expression. Indeed, HEMs expressing OPN3-2 miRNA (the more efficient of the two OPN3-targeted miRNAs) have higher levels of MITF and TYR when compared to HEMs expressing CTRL miRNA (**Fig. 5A**). Conversely, MNT-1 cells expressing OPN3-cMCh had reduced levels of both MITF and TYR, as compared to MNT-1 cells expressing MCh (**Fig. 5B**). These results indicate that OPN3 is a negative regulator of MC1R-meditated melanogenic signaling in human melanocytes.

**Figure 5.**
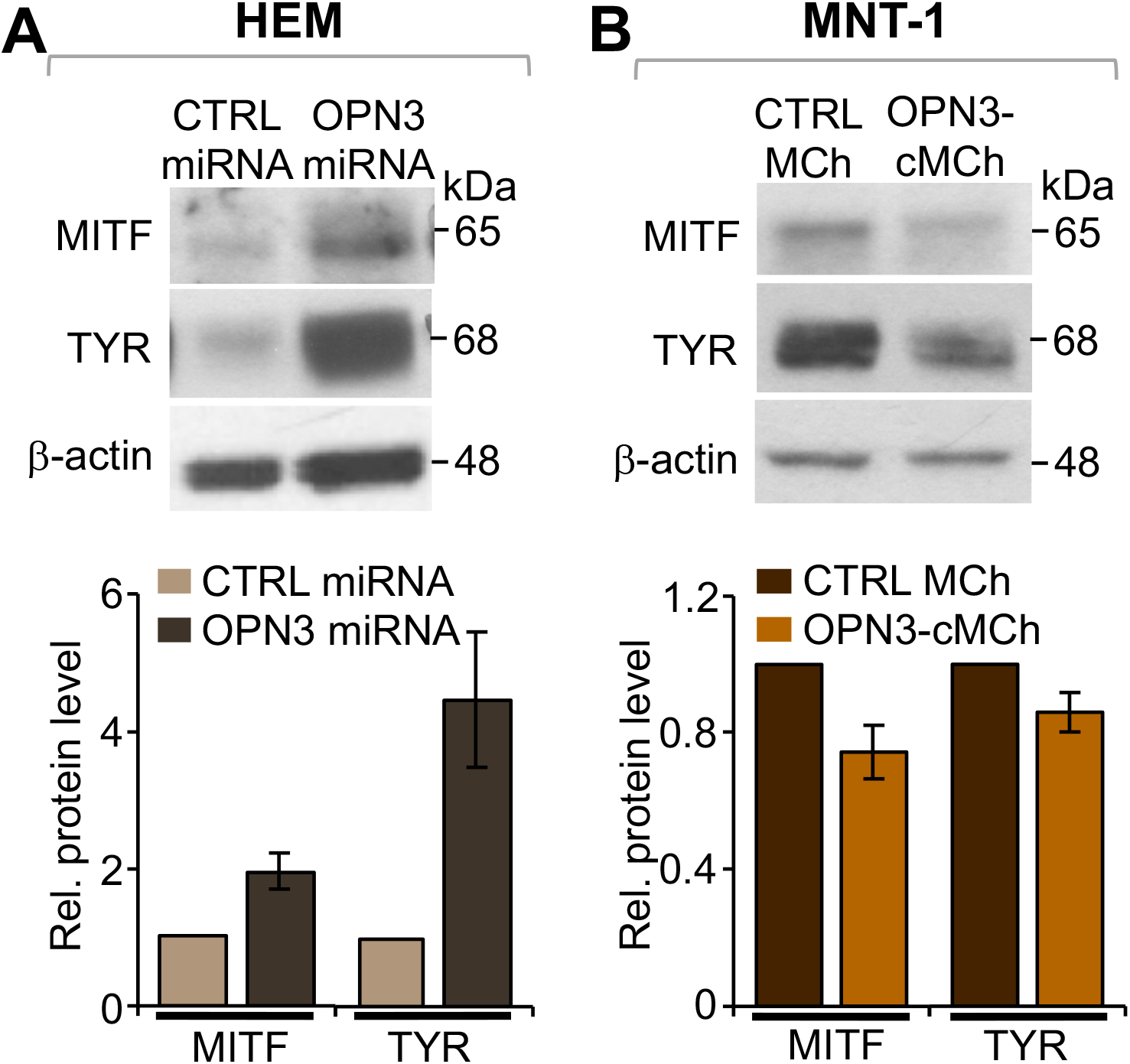
OPN3 modulates MITF and TYR expression. **A. Reduced expression of OPN3 leads to increased MITF and TYR protein levels in HEMs.** Representative western blots of HEMs expressing CTRL or OPN3-targeted miRNA show increased MITF and TYR expression in cells with reduced levels of OPN3 as compared to CTRL. MITF or TYR protein levels measured relative to β-actin, were ~2 fold higher for MITF and ~5 fold higher TYR in HEMs with reduced OPN3 expression, compared to CTRL miRNA expressing cells. n=3 independent experiments for each condition, ± SEM. **B. Increased expression of OPN3 leads to reduced MITF and TYR levels in MNT-1 cells.** Representative western blots of MNT-1 cells stably expressing MCh (CTRL) or OPN3-cMCh show decreased MITF and TYR expression in OPN3-cMCh expressing cells, compared CTRL. MITF and TYR levels measured relative to β-actin were reduced by ~30% for MITF and by ~20% for TYR in cells expressing OPN3-cMCh, as compared CTRL. n=3 independent experiments for each condition, ± SEM.

### MC1R and OPN3 form a complex

Because OPN3 specifically modulates the signaling of MC1R, we proposed that OPN3 and MC1R could form a molecular complex. This would explain why OPN3 and not EP2 modulates MC1R cAMP signaling despite OPN3 and EP2 both being Gαi-coupled: OPN3, MC1R and EP2 are all expressed on the plasma membrane, but MC1R and OPN3 could be localized within the same microdomains.

Several recent reports indicate that GPCRs can form functional and physical interactions [37, 38]. Members of the opsin family, in particular, have been recently shown to form hetero- and homomeric complexes [39, 40]. To determine whether OPN3 and MC1R colocalize to the same cellular compartments, we expressed tagged OPN3 and MC1R in HeLa cells. OPN3-cYFP localized to the plasma membrane, as well as intracellular structures that have partial overlap with the Rab11 marker for recycling endosomes (**Supp. Fig. 4**). To determine if OPN3 colocalization with MC1R is specific to MC1R, we measured OPN3 colocalization with the EP2 receptor, which is also Gαs-coupled. We coexpressed in HeLa cells OPN3-cYFP and either N-terminal HA-tagged MC1R (MC1R-nHA) or N-terminal HA-tagged EP2 receptor (EP2-nHA), immunostained with an anti-HA antibody, and quantified colocalization of YFP and HA. Interestingly, MC1R-nHA and OPN3-cYFP significantly colocalized (>50%) in intracellular structures, while EP2-nHA, localized primarily to the plasma membrane, had <10% overlap with OPN3-cYFP (**Fig. 6A**).

**Figure 6.**
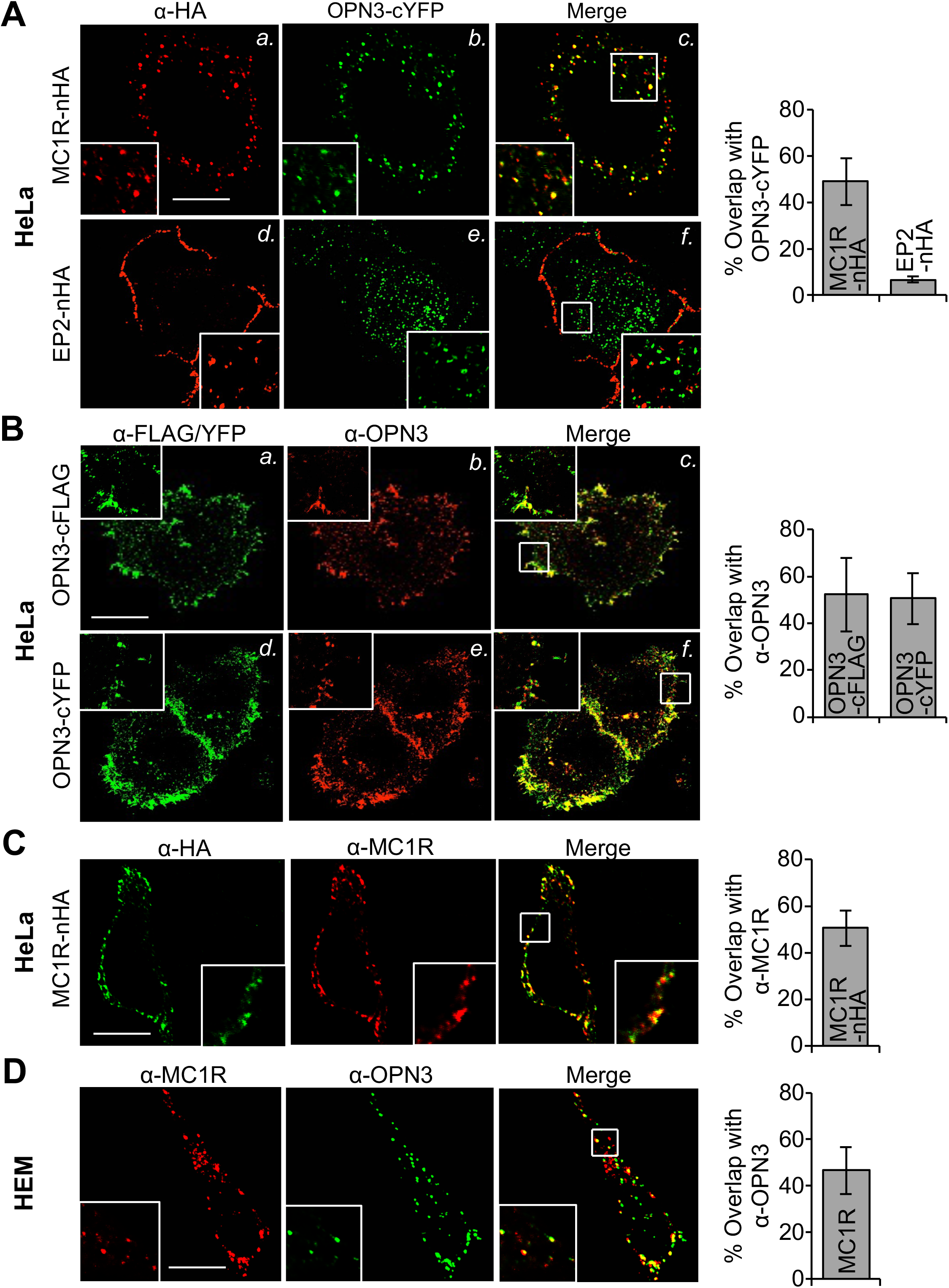
OPN3 and MC1R are localized to the same cellular compartments. **A. OPN3-cYFP is colocalized with MC1R, but not with the EP2 receptor.** Representative fluorescence confocal images of HeLa cells coexpressing OPN3-cYFP and either MC1R-nHA or EP2-nHA and immunostained with anti-HA antibody. OPN3-cYFP shows significant colocalization with MC1R-nHA, but not with the EP2-nHA. Bar graph: the percent overlap between the fluorescent signals of OPN3-cYFP and MC1R-nHA was ~50%, compared to less than 10% for EP2-nHA. n=20 cells from 3 independent experiments, ± SEM. **B. The anti-OPN3 antibody specifically recognizes OPN3-cFLAG and OPN3-cYFP.** Representative fluorescence confocal images of HeLa cells expressing either OPN3-cFLAG or OPN3-cYFP and immunostained with anti-FLAG and anti-OPN3 antibodies. Bar graph: the percent overlap between anti-OPN3 fluorescence and anti-FLAG or YFP fluorescent signal was ~50%, suggesting significant colocalization of anti-OPN3 with both anti-FLAG and YFP signals. n=20 cells from 3 independent experiments, ± SEM. **C. The anti-MC1R antibody specifically recognizes MC1R-nHA.** Representative fluorescence confocal images of HeLa cells expressing MC1R-nHA and co-immunostained with anti-HA and anti-MC1R antibodies. Bar graph: the percent overlap between anti-MC1R and anti-HA fluorescent signals was >50%, indicating significant colocalization. n=25 cells from 3 independent experiments, ± SEM. **D. Endogenously expressed OPN3 and MC1R colocalize in HEMs.** Representative fluorescence confocal images of HEMs co-immunostained with anti-OPN3 and anti-MC1R antibodies. Bar graph: the percent overlap between anti-OPN3 and anti-MC1R fluorescent signals was ~50%, indicating significant colocalization. n=20 cells from 3 independent experiments, ± SEM. Calibration bar: 10 μm.

To confirm that endogenous OPN3 and MC1R colocalize in HEMs, we first validated anti-OPN3 and anti-MC1R antibodies. Cells expressing OPN3-cYFP or OPN3-cFLAG had significant (>50%) overlap between the fluorescent signals obtained with anti-OPN3 or with the respective tags (YFP or FLAG) (**Fig. 6B**). Similarly, cells expressing MC1R-nHA had significant (>50%) overlap between the fluorescent signals of anti-MC1R and anti-HA antibodies (**Fig. 6C**), suggesting that both anti-OPN3 and anti-MC1R antibodies are specific. Immunostaining HEMs with these antibodies revealed that endogenous OPN3 and MC1R are present both at the plasma membrane and in intracellular compartments and exhibit significant colocalization (**Fig. 6D**).

We next determined if OPN3 and MC1R form a physical complex. We first tested if expressed OPN3-cFLAG and MC1R-nHA interact in HeLa cells. Immunoprecipitation with an anti-FLAG antibody and immunoblot with an anti-HA antibody revealed a band corresponding to the molecular weight of MC1R-nHA, but only when both receptors were expressed (**Fig. 7A**). Similarly, immunoprecipitation with anti-HA antibody and immunoblot with anti-FLAG antibody revealed a band corresponding to OPN3-cFLAG, but only when both receptors were expressed (**Supp. Fig. 5**). These results suggest that in HeLa cells OPN3-cFLAG and MC1R-nHA form a complex. To confirm that endogenous OPN3 and MC1R also interact in HEMs, we confirmed that the anti-MC1R antibody detects the same bands as the anti-HA antibody in HeLa cells expressing MC1R-nHA (**Fig. 7B**, left panel). When HEM lysate is immunoprecipitated with anti-OPN3 antibody and immunobotted with the anti-MC1R antibody, a band corresponding to the molecular weight of MC1R is detected (**Fig. 7B**, right panel), indicating that OPN3 and MC1R form a complex in melanocytes.

**Figure 7.**
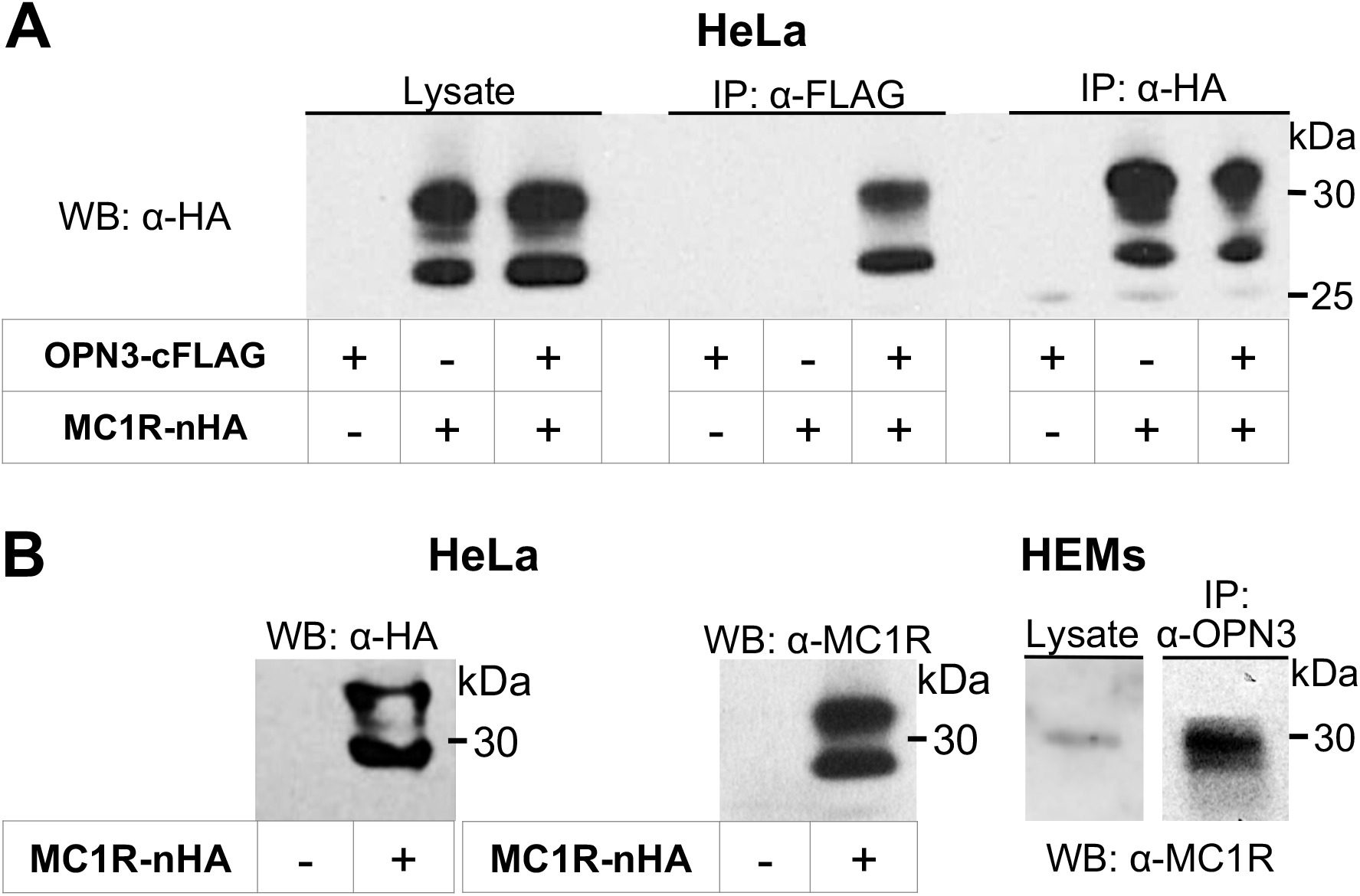
OPN3 and MC1R form a molecular complex. **A. OPN3-cFLAG and MC1R-nHA co-immunoprecipitate.** HeLa cells expressing OPN3-cFLAG, MC1R-nHA, or both, were immunoprecipitated with anti-FLAG and immunoblotted with anti-HA antibodies. A band corresponding to MC1R in the cell lysates (left panel) was detected in the anti-FLAG IP only when both OPN3 and MC1R were expressed (middle panel). The same band was detected by immunoprecipitation with anti-HA antibodies (right panel). **B. Endogenously expressed OPN3 and MC1R co-immunoprecipitate.** Western blot analysis of HeLa cells expressing MC1R-nHA and immunoblotted with anti-MC1R antibody reveal the same size and pattern of bands as detected with the anti-HA antibody in **A** (left panel). HEM lysates co-immunoprecipitated with anti-OPN3 and immunoblotted with anti-MC1R antibodies reveal the same band corresponding to MC1R, suggesting that endogenously expressed OPN3 and MC1R form a complex in HEMs. Representative of n=3 independent experiments for each condition.

Taken together, our results reveal a novel molecular mechanism by which Gαi-coupled OPN3 negatively regulates the cAMP signal resulting specifically from the αMSH-induced activation of MC1R. In addition to the functional interaction, OPN3 and MC1R also form a physical complex, and colocalize at the plasma membrane and also in intracellular structures. The reduced MC1R-mediated cAMP production in the presence of OPN3 leads to reduced activation of MITF, reduced levels of TYR, which ultimately results in decreased melanin production in melanocytes (**Fig. 8**).

**Figure 8.**
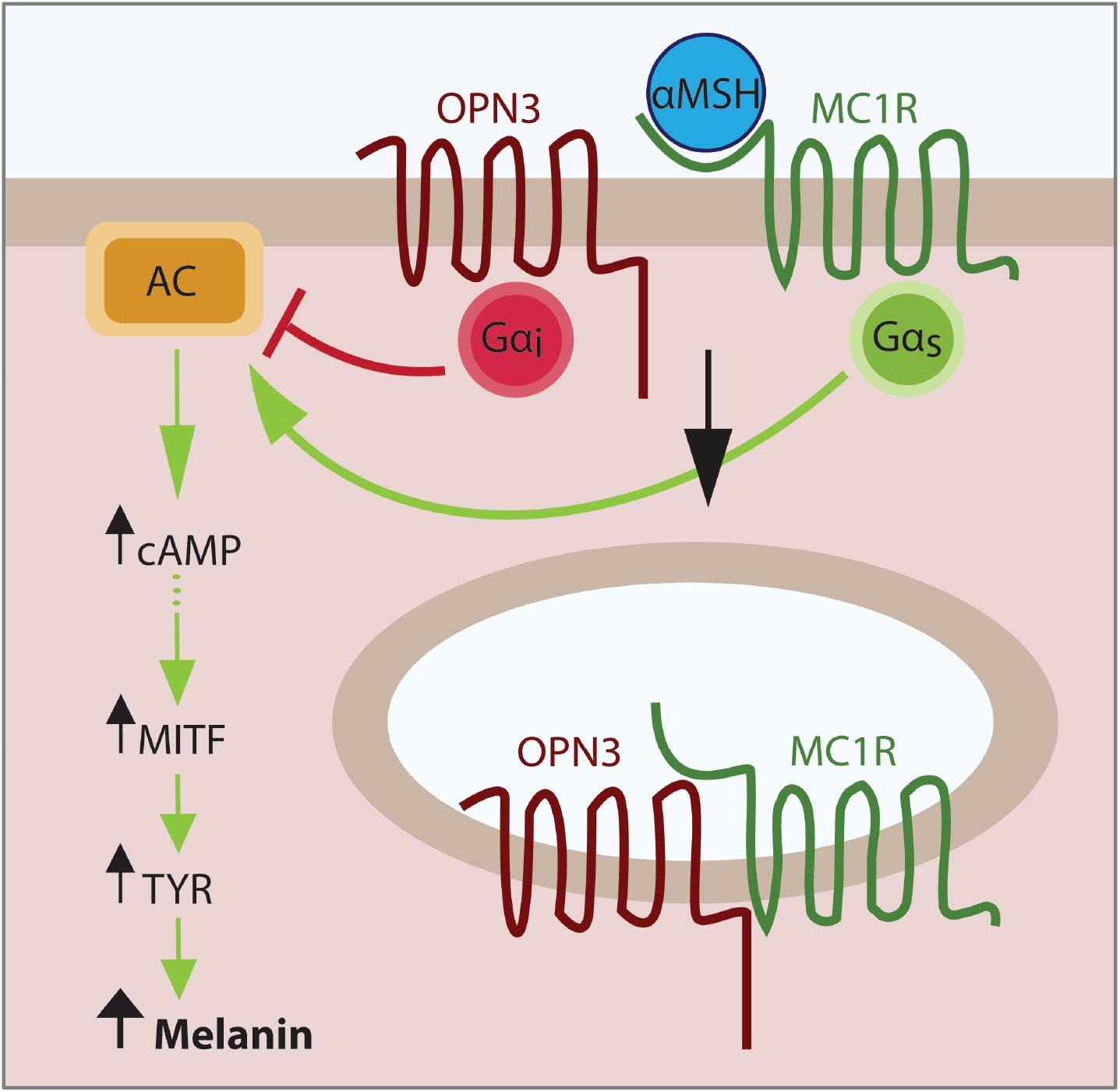
OPN3 and MC1R interact functionally and physically to regulate melanin levels in melanocytes. In melanocytes, αMSH binds to and activates the Gαs-coupled MC1R, leading to stimulation of adenylyl cyclase (AC) and a transient increase in cAMP levels, which, through a series of molecular steps, leads to activation and upregulation of the microphthalmia transcription factor (MITF). MITF controls the expression of tyrosinase (TYR), the main melanogenic enzyme; higher MITF levels will result in more TYR that will generate a higher amount of melanin. Our data suggest that OPN3, via Gαi activation, reduces the amount of cAMP generated by MC1R activation, causing decreased transcription of MITF and, in consequence, of TYR. OPN3 and MC1R form a complex and OPN3 may additionally alter the function of MC1R by enhancing its internalization and lowering the number of receptors available to bind αMSH at the plasma membrane.

## Discussion

In this study we have determined the cellular function of endogenously expressed OPN3 in human melanocytes. We initially hypothesized that OPN3 functioned as the UVR photoreceptor in the UVR-Ca^2+^ melanin signaling pathway previously characterized by our laboratory [14–16]. Surprisingly, our results disproved this hypothesis: we showed that reducing the mRNA and protein levels of OPN3 in melanocytes (**Figs. 1B-D, Supp. Fig. 1B**) did not alter their Ca^2+^-mediated responses to physiological levels of UVR, blue, or green light in melanocytes did not alter the response to UVR, blue or green light (**Figs. 1E and 1F; Supp. Figs. 1D and 1E**). Moreover, when we tested whether purified human OPN3 can respond to visible light, similar to its homologues [24, 25], we found no evidence of visible light absorption in the OPN3 UV-VIS spectrum (**Fig. 2, Supp. Fig. 2B**), despite weakly binding both 11-*cis* and all-*trans* retinal (**Fig. 2C and Supp. Fig. 2A**).

We were puzzled by the finding that human OPN3, despite being able to bind retinal, does not have an absorption peak in the visible range (**Fig. 2C and Supp. Fig. 2A**). This is particularly intriguing considering that OPN3 homologues absorb light in the visible spectrum [24–26]. In addition, unlike two recent reports suggesting that human OPN3 functions in a blue light-dependent manner in human skin [41, 42] and in rat and human pulmonary vasorelaxation [41, 42], we did not detect OPN3 sensitivity to blue light by either spectrophotometric analysis or by monitoring Ca^2+^ levels in melanocytes. One possible scenario is that OPN3 absorbs light outside the range covered by conventional UV-visible spectroscopy (200-700 nm) or that its absorption overlaps with protein absorption (λ_max_=280 nm) and is masked by the large protein peak [43]. Alternatively, OPN3 may be light-insensitive and function in a similar manner as retinal G protein-coupled receptor (RGR), an opsin receptor that binds retinal independent of light stimulus. It was shown that RGR, independent of light, accelerates the conversion of retinyl esters to 11-*cis*-retinal by modulating isomerohydrolase activity [44]. Thus, it is conceivable that OPN3 is not light-sensitive and retinal binding serves another purpose in OPN3 folding, trafficking, or signaling. Future studies are needed to characterize human OPN3 photosensitivity and to understand the role of retinal in OPN3 function.

Reducing or eliminating OPN3 expression in melanocytes led to a surprising observation: these melanocytes had a significantly higher melanin content (**Fig. 3A; Supp. Fig. 1C**). Because melanin production in melanocytes is controlled by the Gαs-coupled MC1R which signals via cAMP, we used genetically encoded cAMP indicators to measure the effect of OPN3 on MC1R-mediated signaling. We found that OPN3 inhibits the MC1R-mediated cAMP response in a Gαi–dependent manner (**Fig. 4**), indicating that MC1R and OPN3 functionally interact. The functional interaction between OPN3 and MC1R is similar to the mechanism described for the cross-regulation of the G protein-coupled estrogen receptor (GPER) and steroid hormone adipoQ receptor 7 (PAQR7), in which PAQR7 regulates skin pigmentation through Gαi-mediated downregulation of GPER activity [45]. The ability of OPN3 to negatively regulate MC1R signaling does not appear to require retinal binding to OPN3, as the OPN3(K299G) mutant is able to modulate MC1R-mediated signaling, despite not binding retinal (**Fig. 4**), further supporting the idea that OPN3 functions in a light-independent manner in melanocytes.

Human OPN3 is likely to be coupled to Gαi, similar to its homologues [24–26]. Indeed, using PTX to prevent Gαi activation eliminated the inhibitory effect of OPN3 on MC1R-mediated cAMP signaling (**Figs. 4Aiii and 4Aiv**). These results indicate that OPN3 is coupled to Gαi either constitutively or, alternatively, light might turn off OPN3 that is coupled to Gαi. In this case, exposure to light would uncouple OPN3 from Gαi, resulting in an increase in the baseline cAMP. Initially, we were not able to measure light-induced OPN3-dependent changes in cAMP in melanocytes, because the FRET indicator Epac H187, used to monitor cAMP, requires illumination of the cells with visible light in the blue-green range (~ 400– 530 nm). Thus, if the light used for Epac H187 imaging was sufficient to inactivate OPN3, the increase in cAMP could occur prior to starting image acquisition. We circumvented this problem by using a recently developed genetically encoded red cAMP indicator, R-FlincA (λ_ex_ = 587 nm) [34]. To determine if blue, green or UV radiation elicits an OPN3-dependent change in baseline cAMP levels, we used MNT-1 cells (with already low endogenous levels of OPN3 (**Fig 3B**)) to express R-FlincA together with either OPN3-cYFP or YFP. We monitored R-FlincA fluorescence emission before, during, and after stimulation with 200 mW/cm^2^ of blue, green or UV radiation (**Supp. Fig. 6**). We did not measure any light-induced or OPN3-dependent changes in cAMP levels, suggesting that the amount of UVR, blue, or green light applied does not modulate OPN3 activity. This would suggest that OPN3 is more likely to have full or partial constitutive activity. The constitutive activity of OPN3 might require conformational changes induced through direct or indirect interaction with MC1R. In fact, MC1R has unusually high partial constitutive activity, which is responsible for the baseline pigmentation in our skin and hair. The constitutive activity of OPN3, coupled to Gαi, could be responsible for regulating both baseline cellular cAMP levels in the absence of MC1R agonists and cAMP levels upon MC1R activation.

Our colocalization (**Fig. 6**) and co-immunoprecipitation (**Fig. 7**) data indicate that OPN3 and MC1R reside in the same microdomains where they form a molecular complex. The presence of both OPN3 and MC1R in the same microdomains could explain the specific OPN3-mediated regulation of MC1R, but not of EP2 receptor signaling (**Fig. 6**). The microdomain localization might allow OPN3 to become activated by binding to MC1R directly or via other proteins in the complex. Alternatively, the microdomain might contain the specific adenylyl cyclase that is activated by MC1R and inhibited by OPN3, or a particular phosphodiesterase that could be modulated by OPN3. In all these scenarios, OPN3 would not be able to modulate signaling via cAMP evoked by Gαs-coupled receptors that are not localized to the same microdomains, like EP2 receptors.

Heteromeric receptors have been described for other GPCRs and have been shown to have altered ligand binding, signaling, and internalization properties compared to the individual GPCRs [37, 38]. For example, in the brain, the melanocortin 3 receptor (MC3R), of the same melanocortin family as MC1R, and the ghrelin receptor (GHSR) form heteromers with altered melanocortin- and ghrelin-induced intracellular responses to regulate energy metabolism [46, 47]. Interestingly, OPN3 was initially identified in deep regions of the brain (hence the initial name “encephalopsin”) [21] including the hypothalamus, where MC3R is also expressed [48–50]. This raises the intriguing question of whether OPN3 functions in the brain similar to how it does in melanocytes: could OPN3 interact and negatively regulate the function of other melanocortin receptors like MC3R? OPN3 is not the only opsin that functions as part of a dimeric complex; a functional homomeric complex has also been reported for rhodopsin (OPN2) and cone opsin (OPN1) [40]. These complexes are mediated by residues within the fifth transmembrane domain of human red and green opsins [39], and by transmembrane domain one and helix eight for rhodopsin dimerization [40]. Whether OPN3 and MC1R directly interact through domain coupling or are part of a larger complex was beyond the scope of our study, but will be determined by future studies.

Our immunostaining results of both expressed and endogenous OPN3 and MC1R show that the two receptors colocalize at the plasma membrane as well as in intracellular compartments. Because recent studies showed that activated GPCRs are internalized and could continue to signal in endosomal compartments [51], it is intriguing to hypothesize that MC1R and OPN3 continue to signal even when they are no longer at the plasma membrane. We wondered if the colocalization of OPN3 and MC1R in melanocytes depends on the activation state of MC1R. We compared the localization of OPN3 and MC1R in growth media not containing αMSH and 1, 3 or 6 h after αMSH stimulation (**Supp. Fig. 7**). We found that αMSH-mediated activation of MC1R did not cause a significant shift in the fraction of MC1R at the plasma membrane vs. intracellular organelles and, implicitly, did not alter the fraction of MC1R colocalized with OPN3, suggesting that both active and inactive forms of MC1R are associated with OPN3.

The findings presented here expand our understanding of OPN3 function and its role as an extraocular opsin in human skin. We have uncovered a novel light-independent function of OPN3 in regulating melanin levels in human melanocytes. Taking into consideration that retinoic acid, a derivative of vitamin A, is widely used in skin treatments such as hyperpigmentation [52–54] and that OPN3 can bind retinal, OPN3 could yield a novel therapeutic target for skin pigmentation disorders such as melasma, and the OPN3-MC1R complex may reveal novel molecular mechanisms for opsin function and for regulating melanin production in melanocytes.

## Materials and Methods

### Cell culture and transfection

All cell culture reagents were purchased from ThermoFisher Scientific, unless otherwise stated.

<u>HeLa cells</u>, used for immunostaining and biochemical analyses of expressed tagged OPN3 and MC1R, were maintained at 37°C and 5% CO_2_ in DMEM supplemented with 5% FBS and 100 units/ml penicillin/streptomycin and transiently transfected using Lipofectamine 2000 according to manufacturer’s recommendations.

<u>HEK293-GnTl^−^ cells</u>, used for expression and purification of OPN3 variants and OPN2 (as a control) for UV-visible spectroscopy, were maintained under standard conditions in DMEM supplemented with 10% FBS and 100 units/ml penicillin/streptomycin and transfected using calcium phosphate precipitation, as previously described [55].

We used three types of melanocytes:

▫ <u>Normal primary neonatal human epidermal melanocytes (HEMs)</u>. HEMs lines derived from at least three individuals were purchased and maintained under standard conditions in Medium 254 supplemented with human melanocyte growth supplement (HMGS-2) and 100 units/ml penicillin/streptomycin. For miRNA experiments, HEMs were transduced with either OPN3-targeted or control miRNAs using BLOCK-IT™ Lentiviral RNAi expression system according to manufacturer’s protocol. The lentiviral transduction rates were ~60% as detected by coexpression of MCherry. HEMs expressing miRNAs were selected with blasticidin (4 μg/ml) for at least 14 days.
▫ <u>Immortalized normal human epidermal melanocytes (Hermes 2b)</u>, purchased from Cell Bank, UK (https://www.sgul.ac.uk/depts/anatomy/pages/Dot/Cell%20bank%20holdings.htm#Hermes) are HEMs immortalized with hTERT (puro vector) only and maintained at 37°C and 10% CO_2_ in RPMI 1640 media supplemented with 20% FBS, 200 nM TPA, 200 pM cholera toxin, 10 ng/ml human stem cell factor, 10 nM endothelin and 100 units/ml penicillin/streptomycin. Hermes 2b cells were transiently transfected using Nucleofector™ Kits for Human Melanocytes (Lonza) according to manufacturer’s instructions. For CRISPR/Cas9 knock-out experiments, Hermes 2b cells were transduced with OPN3-targeted lentiCRISPR v2 (single guide RNA: ggccacggctactgggacgg). Transduced Hermes 2b cells were selected with puromycin (10 μg/ml) and clonal lines were isolated and maintained.
▫ <u>Human pigmented melanoma melanocytes (MNT-1)</u> were maintained under standard conditions in DMEM supplemented with 18% FBS, 10% AIM-V and 100 units/ml penicillin/ streptomycin and transiently transfected using Nucleofector™ Kits for Human Melanocytes (Lonza) or magnetofection with PolyMag Neo magnetic beads (OZ Biosciences), according to manufacturer’s instructions.

### DNA constructs

cDNA encoding full length human OPN3 was obtained by RT-PCR using RNA extracted from HEMs. Different OPN3 variants were cloned as summarized in the table below. Mutations were introduced by site-directed mutagenesis using the QuickChange^®^ Site-Directed Mutagenesis Kit (Stratagene). The human MC1R-n(3xHA) and EP2-n(3xHA) in pcDNA3.1 expression vectors were purchased from www.cDNA.org. EEA1-nRFP, Rab7-nRFP, Rab11-nDsRed, Rab9-nDsRed, MEM-nDsRed were obtained from www.addgene.org. All constructs were confirmed by sequencing.

| Construct | Vector | Restriction Sites |
| --- | --- | --- |
| OPN3-cMCh | pcDNA4/TO-cMCh | HindIII-XhoI |
| OPN3-cHA | pcDNA4/TO-cHA | EcoRI-XhoI |
| OPN3-nHA | pcDNA4/TO-nGFP | EcoRI-XhoI |
| OPN3-nHA-ΔC | pcDNA4/TO-cGFP | EcoRI-XhoI |
| MCh CTRL | pcDNA4/TO-cMCh | -- |
| OPN3-cFLAG | pcDNA3.1-cFLAG | HindIII-XhoI |
| OPN3-cYFP | pcDNA3.1-cYFP | BamHI-XhoI |
| OPN3(K299G)-cYFP | pcDNA3.1 C-YFP | BamHI-XhoI |
| YFP CTRL | pcDNA3.1-cYFP | -- |
| miRNA OPN3-1 and 2 | pLenti V5-DEST | Attb1-Attb2 |
| OPN3-c1D4 | pMT3 | EcoRI-NotI |
| OPN3ΔC-c1D4 | pMT3 | EcoRI-NotI |
| OPN3(K299G)ΔC-c1D4 | pMT3 | EcoRI-NotI |

### Immunoprecipitation and Western blot

Cells were rinsed with cold PBS before addition of 500 μl lysis buffer [300 mM NaCl, 50 mM Tris-HCl (pH 7.4), 1% Triton X-100, and protease inhibitor mix (Roche)]. Cells were scraped and homogenized using a 22G syringe needle. Lysates were rotated end-over-end for 1 h at 4°C then centrifuged at 14,000 rpm for 30 min at 4°C to remove cell debris. 15 μl of 50% (w/v) protein A/G or protein A beads (Santa Cruz Biotechnology) were added to the supernatant and rotated for 30 min to preclear the samples. Samples were centrifuged at 14,000 rpm for 5 min, and the agarose pellet was discarded. Samples were split into two aliquots and mixed with 25 μl of primary antibody conjugated to protein A or protein A/G beads and rotated overnight at 4°C. Immunoprecipitates were collected by centrifugation at 7,000 rpm for 5 s, washed three times with wash buffer [300 mM NaCl, 50 mM Tris-HCl (pH 7.4), and 0.1% Triton X-100], and solubilized with 10 μl elution buffer [100 mM Tris-HCl, 1% SDS, 10 mM DTT] and 5 μl of 4X NuPAGE LDS sample buffer (ThermoFisher Scientific). For Western blots and immunoprecipitation experiments the following primary antibodies were used: anti-HA (Roche, 11867423001), anti-FLAG (Sigma-Aldrich, F1804-200UG), anti-OPN3 (Santa Cruz Biotechnology, sc-98799), anti-OPN3 (ABclonal, A15803), anti-MC1R (Santa Cruz Biotechnology, sc-6875), anti-MITF (ThermoFisher Scientific, MA514146), anti-TYR (Santa Cruz Biotechnology, sc-7833), anti-β-actin (ThermoFisher Scientific, MA515739) and anti-integrin α5 (Santa Cruz Biotechnology, H-104). The primary antibodies were detected by incubation with HRP conjugated goat anti-rat, goat anti-mouse or donkey anti-goat secondary antibodies.

### Immunofluorescence

Cells seeded on glass coverslips were fixed with 4% paraformaldehyde at room temperature for 10 min, incubated with blocking solution [0.2% saponin, 0.1% BSA, 0.02% sodium azide] for 1 h, followed by overnight incubation with primary antibodies (1:200). For visualization, fluorescently labeled secondary antibodies (Life Technologies) (1:1000) were used. Confocal images were acquired using an Olympus FV3000 confocal microscope and analyzed using Image J software. The Pearson correlation coefficient (PCC) was calculated for each cell, as previously described [56]. Two proteins were considered significantly colocalized if the overlap was higher than 50% and not colocalized if was lower than 10% [57]. As a positive control for the colocalization analysis we calculated PCC for two well-characterized lysosomal proteins, TPC2 and LAMP1, expressed in HeLa cells, which showed ~50% overlap (**Supp. Fig. 8**).

### Ca^2+^ imaging and light stimulation

Cells cultured on glass coverslips were incubated for 20 min with 12 μM all-*trans* retinal (Sigma-Aldrich) in Ringer’s solution [150 mM NaCl, 5 mM KCl, 1.8 mM CaCl_2_, 1.2 mM MgCl_2_, 10 mM HEPES, 10 mM D-glucose, pH 7.4], followed by wash and 20 min incubation with 7.5 μM Fluo4-AM (Life Technologies) and 250 μM sulfinpyrazone (uridine 5’-diphospho-glucuronosyltransferase, Sigma-Aldrich) in Ringer’s solution. Ca^2+^ imaging was performed using an inverted microscope (Olympus IX71). Sequential images were acquired with a 20x objective every 2 s before, during and after light stimulation: 200 mJ/cm^2^ UVR (λ_max_=360 nm); 200 mJ/cm^2^ blue (λ_max_=460 nm) or green (λ_max_=560 nm) light. UVR was applied by using a 400 nm short pass and 280 nm long pass filters (Newport) attached to a 200 W Hg-Xe arc lamp (Newport) as previously described [16]. For blue and green light, 460 and 560 nm LED light sources (Prizmatix) were used. For all experiments, cells were exposed to 20 mW/cm^2^ radiation for 10 seconds, resulting in a total dose of 200 mJ/cm^2^. Ionomycin (1 μM) was added at the end of each experiment to elicit a maximal Ca^2+^response, used for normalization. Changes in fluorescence intensity of individual cells as a function of time were obtained using MetaMorph software, then analyzed with MatLab and Microsoft Excel.

### cAMP imaging

The genetically encoded cAMP indicator mTurq2Del-EPAC(dDEPCD)Q270E-tdcp173Venus(d) EPAC-S^H187^ (Epac H187) (K_d_=4.0 μM) and red fluorescent cAMP indicator R-FlincA (K_d_=0.3 μM) were generous gifts from the Jalink Laboratory (Netherlands Cancer Institute) and Horikawa Laboratory (Tokushima University), respectively. MNT-1 cells were transfected with Epac H187 and ~24 h after transfection cells were serum-starved in OPTI-MEM (ThermoFisher Scientific) for another ~24 h. Hermes 2b cells were transfected with R-FlincA and ~6 h after transfection cells were serum-starved for another ~6 h. For PTX treatment, serum starved cells were incubated with 200 ng/ml PTX for 4 h before the experiment. Coverslips were transferred to an imaging chamber with Ringer’s solution. For Epac H187 imaging, sequential fluorescence images were acquired with MetaMorph software on an inverted microscope every 10 s using CFP and FRET filter cubes: λ_ex_ = 430 nm and CFP and YFP emissions were detected simultaneously using 470±20 nm and 530±25 nm band-pass filters. For R-FlincA imaging, images were acquired every 10 s using an MCherry filter cube (λ_ex_ = 587 nm). For both Epac H187 and R-FlincA, after acquiring 18 baseline images (3 min), 1 μM NDP-αMSH (Sigma-Aldrich) or 5 μM prostaglandin (Sigma-Aldrich) were added. After 54 images (9 min), a mix of 25 μM forskolin (FSK, Sigma-Aldrich) and 100 μM 3-isobutyl-1-methylxanthine (IBMX, Sigma-Aldrich) was added to elicit a maximal cAMP response, used for normalization. For MNT-1 cells expressing R-FlincA, cells were exposed to 200 mJ/cm^2^ UVR (λmax=360 nm), blue (λ_max_=460 nm) or green (λ_max_=560 nm) light after 100 s baseline; after 5 min, FSK+IBMX was added. Fluorescence emission intensities for Epac H187 and R-FlincA were calculated as F=F_CFP_/F_YFP_ and F=F_MCh_, respectively. Normalized fluorescence intensities were quantified using F_norm_(t) = (F_cell_(t)–F_min_)/(F_FSK+IBMX_ – F_min_), where F_cell_ is the fluorescence of an intracellular region of interest, F_FSK+IBMX_ is the maximal fluorescence with FSK and IBMX, and F_min_ is the baseline fluorescence before stimulation. Light-induced changes in fluorescence intensity were quantified using MetaMorph and Excel software (Microsoft). NDP-αMSH, prostaglandin, FSK and IBMX were solubilized in DMSO (Sigma-Aldrich) at >100x the final concentration, so that the final DMSO concentration in the imaging chamber remained < 1% (v/v) for all experiments.

### Melanin assay

Confluent melanocytes cultured in 35 mm dishes were lysed in 1% Triton X100 and centrifuged at 14,000 rpm for 30 min at 4°C to separate melanin from solubilized protein. Melanin pellets were solubilized in 1 M NaOH at 85°C for 1+ h. The volumes of the solubilized protein and solubilized melanin were noted. Spectrophotometric analysis of melanin content was determined by measuring absorbance at 405 nm and using a calibration curve obtained with synthetic melanin, as previously described [16]. Total melanin was determined as the product of the melanin concentration measured spectrophotometrically and the total volume of solubilized melanin. The protein content for each sample was measured using BCA protein assay kit (Pierce™, ThermoFisher Scientific). Total protein was determined as the product of the protein concentration measured with BCA and the volume of solubilized protein. Cellular melanin concentration was determined as total melanin/total protein for each condition; relative melanin content was calculated as the ratio of cellular melanin concentration for each experimental condition to control.

### UV-visible spectroscopy

HEK293-GnTl^−^ cells plated on 100 mm culture dishes were transfected, using calcium phosphate precipitation, with OPN3ΔC-c1D4 or OPN3(K299G)ΔC-c1D4 (the ΔC truncation of amino acids 315-402 at the C-terminus increased protein yield). After three days, cells were harvested, centrifuged at 3,500 rpm for 20 min and re-suspended in PBS. All subsequent steps were performed in the dark. Cells were treated with 4.8 mM all-*trans* or 11-*cis* retinal at 4°C for 30 min, lysed with 1% n-Dodecyl-β-D-maltoside (DDM, Sigma Aldrich) at 4°C for 1 h, then centrifuged at 3,500 rpm for 20 min. The supernatant containing the solubilized protein was incubated with pre-conjugated 1D4 antibody-Sepharose beads at 4°C for 2 h, then run through a disposable plastic column (ThermoFisher Scientific) and washed with 0.1% DDM. Proteins were eluted with 0.4 mM 1D4 peptide solution at 4°C. Absorbance spectra were measured on a Cary 50-UV visible spectrometer between 200 and 800 nm as previously described [55]. To test the presence of a Schiff-base bond between chromophore and the K299 residue of OPN3, 0.8% SDS and 80 μM NH_2_OH were added to create retinal oxime, which absorbs maximally at 360 nm [31].

### Statistical analysis

For each tested condition, several replicate experiments were performed and results were averaged. All data are given as means ± SEM. Statistical differences among the experimental groups were analyzed by two-sided Student t test when comparing two experimental groups. Significance was defined as *p < 0.05 and **p < 0.01.

## Acknowledgements

We thank members of the Oancea Laboratory for technical assistance and insightful discussions. We thank Dr. K. Jalink (Netherlands Cancer Institute) and Dr. K. Horikawa (Tokushima University) for generously providing the cAMP indicators Epac H187 and R-FlincA, respectively. This work was supported by NIAMS R01 AR066318 (E.O.), NIGMS training grant T32 GM077995 (L.E.O.), and Brown University Institute for Brain Science/The Suna and Inan Kirac Foundation fellowship (R.N.O.).

