## Supplemental Figures 1-8 for "The human non-visual opsin OPN3 regulates pigmentation of epidermal melanocytes through interaction with MC1R"

### Supplementary Material

**Supplementary Figures:**

**Supp. Fig. 1 – Supp. Fig. 8**

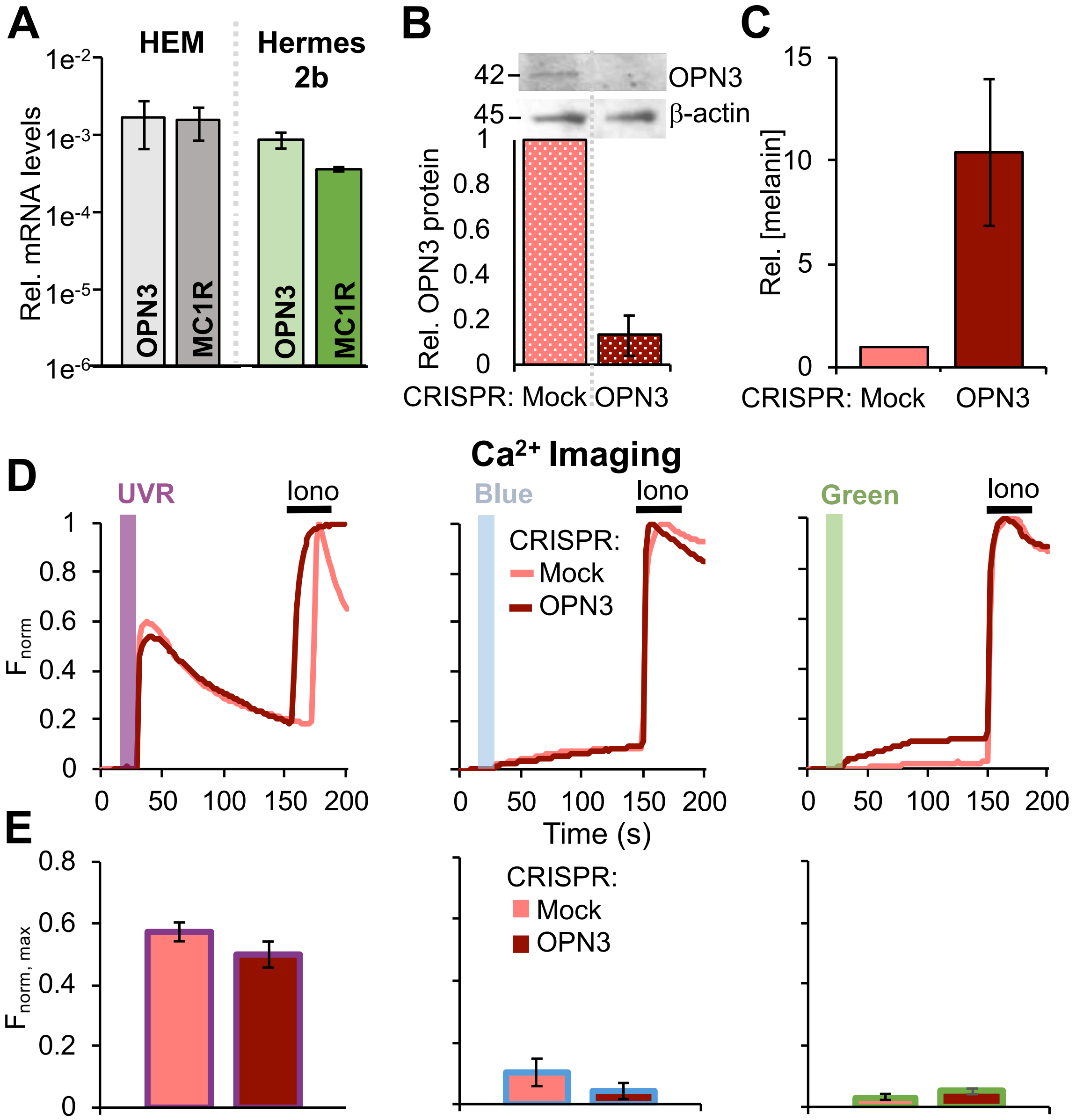


**Supp. Fig. 1. OPN3 does not alter light-induced $Ca^{2+}$ responses in Hermes 2b cells.**

A. **mRNA levels of OPN3 and MC1R in HEMs and Hermes 2b melanocytes,** measured by qPCR analysis. n=3 independent experiment, ± SEM.

B. **Hermes 2b cells treated with OPN3-targetted CRISPR/Cas9 have negligible levels of OPN3 protein.** Representative Western blot of Hermes 2b cells expressing OPN3-targeted CRISPR/Cas9 or were mock-transfected and blotted with anti-OPN3 or with anti-β-actin antibodies. Bar graph represents relative OPN3 protein level. n=2, ± SEM.

C. **Hermes 2b cells lacking OPN3 expression have increased cellular melanin.** Hermes 2b cells expressing OPN3-targetted CRISPR/Cas9 have significantly higher average melanin levels compared to control cells. n=2, ± SEM.

D. **Light-induced $Ca^{2+}$ signaling in Hermes 2b cells is not OPN3-dependent.** Fluorescent $Ca^{2+}$ imaging of Hermes 2b cells expressing endogenous OPN3 (Mock) or lacking OPN3 (CRISPR OPN3) stimulated with 200 mJ/cm$^2$ ultraviolet (UVR, $\lambda_{max}$=360 nm), blue ($\lambda_{max}$=450 nm) or green ($\lambda_{max}$=550 nm) light and normalized to the maximal $Ca^{2+}$ response to ionomycin (Iono). Each trace is a representative of the average of 5-12 cells from one coverslip.

E. **Average amplitude of $Ca^{2+}$ responses of Hermes 2b cells under conditions shown in D.** n=20-35 total cells per condition from 3-4 independent experiments, ± SEM.

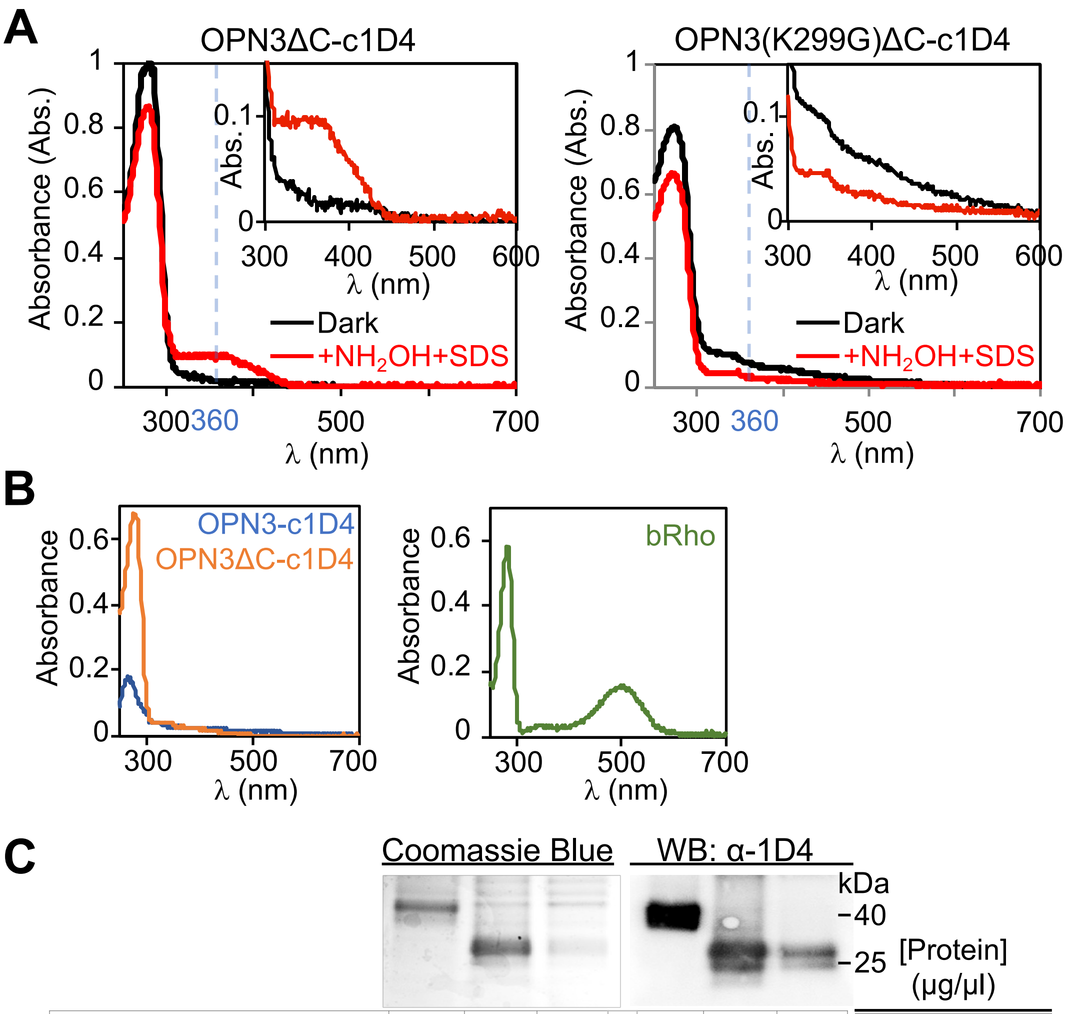


| | | | | | | | |
|---|---|---|---|---|---|---|---|
| OPN3-c1D4 | + | - | - | + | - | - | 0.15 ± 0.04 |
| OPN3ΔC-c1D4 | - | + | - | - | + | - | 0.73 ± 0.04 |
| OPN3(K299G)ΔC-c1D4 | - | - | + | - | - | + | 0.63 ± 0.05 |

**Supp. Fig. 2**. **OPN3, but not OPN3(K299G) binds all-*trans* retinal.**

**A. Absorption spectra of purified OPN3ΔC-c1D4 and OPN3(K299G)ΔC-c1D4** pre-incubated with all-*trans* retinal as measured in the dark (black) and after hydroxylamine and sodium dodecyl sulfate ($NH_2OH$+SDS) treatment (red). After $NH_2OH$+SDS treatment, OPN3ΔC-c1D4—but not OPN3(K299G)ΔC-c1D4—exhibited an absorption peak at $\lambda_{max}$=360 nm, corresponding to retinal oxime. Insets: Similar to Fig. 2C, the retinal oxime peak was ~10 times smaller than the protein peak.

**B. Absorption spectra of purified OPN3-c1D4 (blue), OPN3ΔC-c1D4 (orange) and bovine rhodopsin (bRho, green)** incubated with 11-*cis* retinal and measured in the dark. OPN3ΔC-c1D4 has >3 fold higher protein expression compared to OPN3-c1D4. bRho exhibits the expected absorption peak at $\lambda_{max}$=500 nm.

**C. Purified protein samples of OPN3-c1D4, OPN3ΔC-c1D4 and OPN3(K299G)ΔC-c1D4.** OPN3 variants were expressed in HEK293 GnTI$^-$ cells, purified, run on SDS-PAGE, then stained with Coomassie Blue or immunoblotted with anti-1D4 antibody. The band corresponding to OPN3 in the anti-1D4 Western blot (WB) was also the main band detected by Coomassie Blue staining. Representative of n=2 independent experiments.

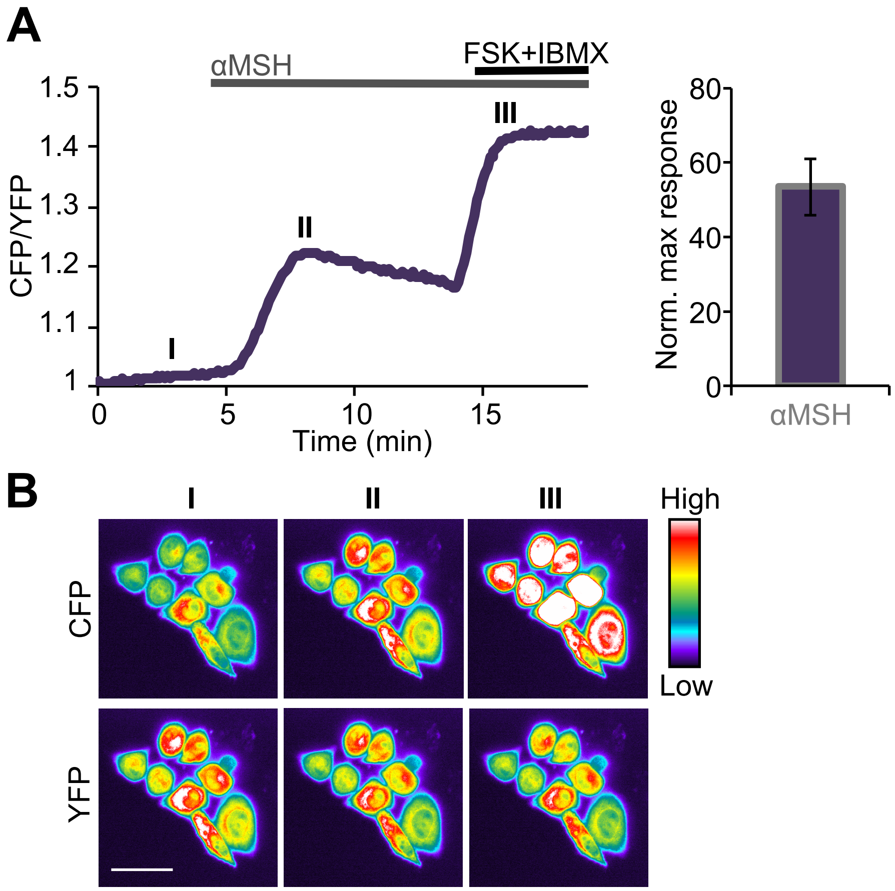


**Supp. Fig. 3. αMSH stimulation of HeLa cells expressing MC1R-nHA leads to a significant increase in intracellular cAMP, as measured with Epac H187.**

**A. Representative average trace of HeLa cells expressing MC1R-nHA and Epac H187.** Stimulation with αMSH led to a rapid increase in cAMP levels, measured by the ratio of CFP/YFP intensity of Epac H187. The bar graph represents the average amplitude of the αMSH-induced cAMP responses normalized to the maximal response obtained with FSK+IBMX. n=3 independent experiments, ± SEM.

**B. Pseudo-color images of HeLa cells acquired during the experiment shown in A.** Times indicated on the graph: baseline (I), after αMSH (II) and after FSK+IBMX (III) stimulation. Calibration bar, 20 μm.

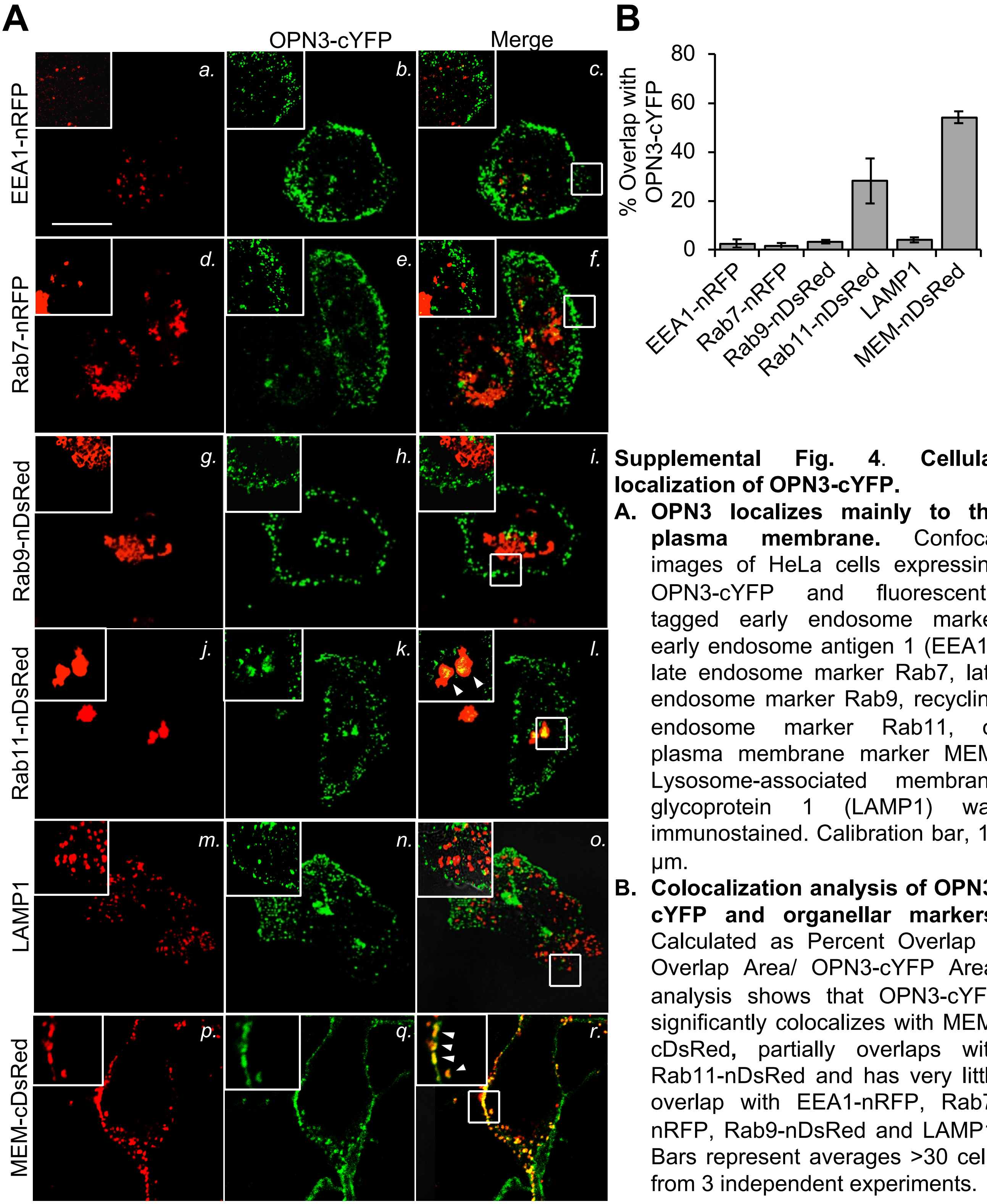


**Supplemental Fig. 4. Cellular localization of OPN3-cYFP.**

A. **OPN3 localizes mainly to the plasma membrane.** Confocal images of HeLa cells expressing OPN3-cYFP and fluorescently tagged early endosome marker early endosome antigen 1 (EEA1), late endosome marker Rab7, late endosome marker Rab9, recycling endosome marker Rab11, or plasma membrane marker MEM. Lysosome-associated membrane glycoprotein 1 (LAMP1) was immunostained. Calibration bar, 10 µm.

B. **Colocalization analysis of OPN3-cYFP and organellar markers.** Calculated as Percent Overlap = Overlap Area/ OPN3-cYFP Area, analysis shows that OPN3-cYFP significantly colocalizes with MEM-cDsRed, partially overlaps with Rab11-nDsRed and has very little overlap with EEA1-nRFP, Rab7-nRFP, Rab9-nDsRed and LAMP1. Bars represent averages >30 cells from 3 independent experiments.

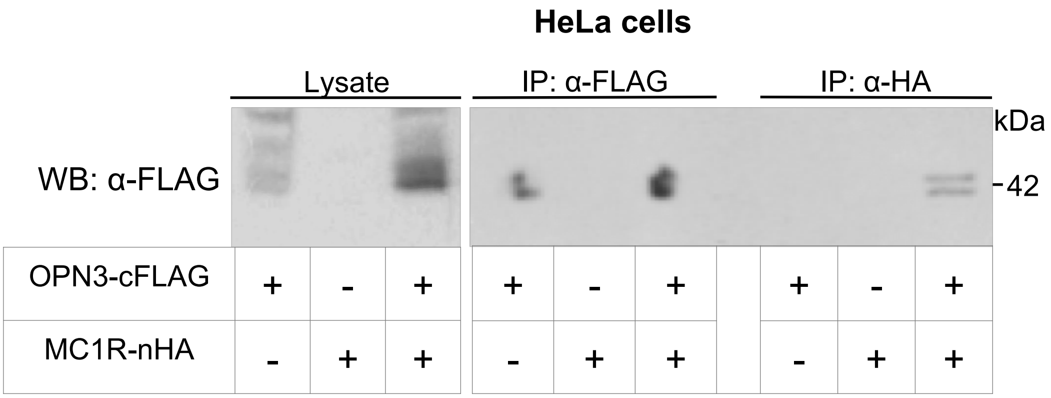


**Supp. Fig. 5**. **OPN3 and MC1R coimmunoprecipitate in HeLa cells.** HeLa cells expressing OPN3-cFLAG, MC1R-nHA, or both were immunoprecipitated (IP) with anti-FLAG or anti-HA antibodies, followed by anti-FLAG blotting. Bands at ~42 kDa correspond to positive staining for OPN3-cFLAG. Bands corresponding to the anti-FLAG IPs indicate that the FLAG antibody immunoprecipitates OPN3-cFLAG. Bands corresponding to the anti-HA IPs indicate that OPN3-cFLAG and MC1R-nHA coimmunoprecipitate.

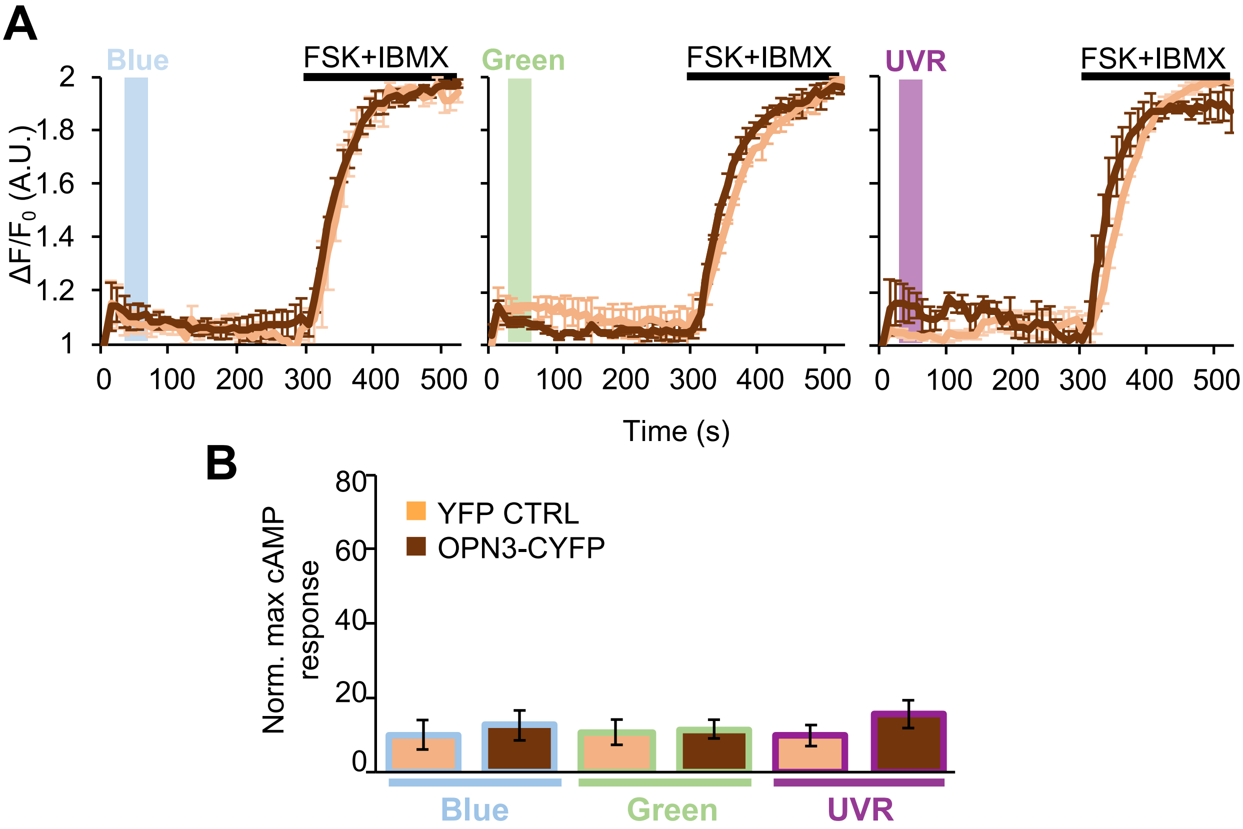


**Supp. Fig. 6. OPN3 does not mediate light-induced cAMP responses in MNT-1 cells.**

A. **MNT-1 cells expressing RFlincA and OPN3-cYFP or YFP alone (CTRL) and stimulated with 200 mW/cm² of blue, green, or UV light did not elicit significant cAMP responses.** n= 5-13 cells per condition, ± SEM.

B. **The average normalized amplitudes of blue, green, or UV light-induced cAMP responses for all conditions in A.** n = 2-3 independent experiments per condition, ± SEM.

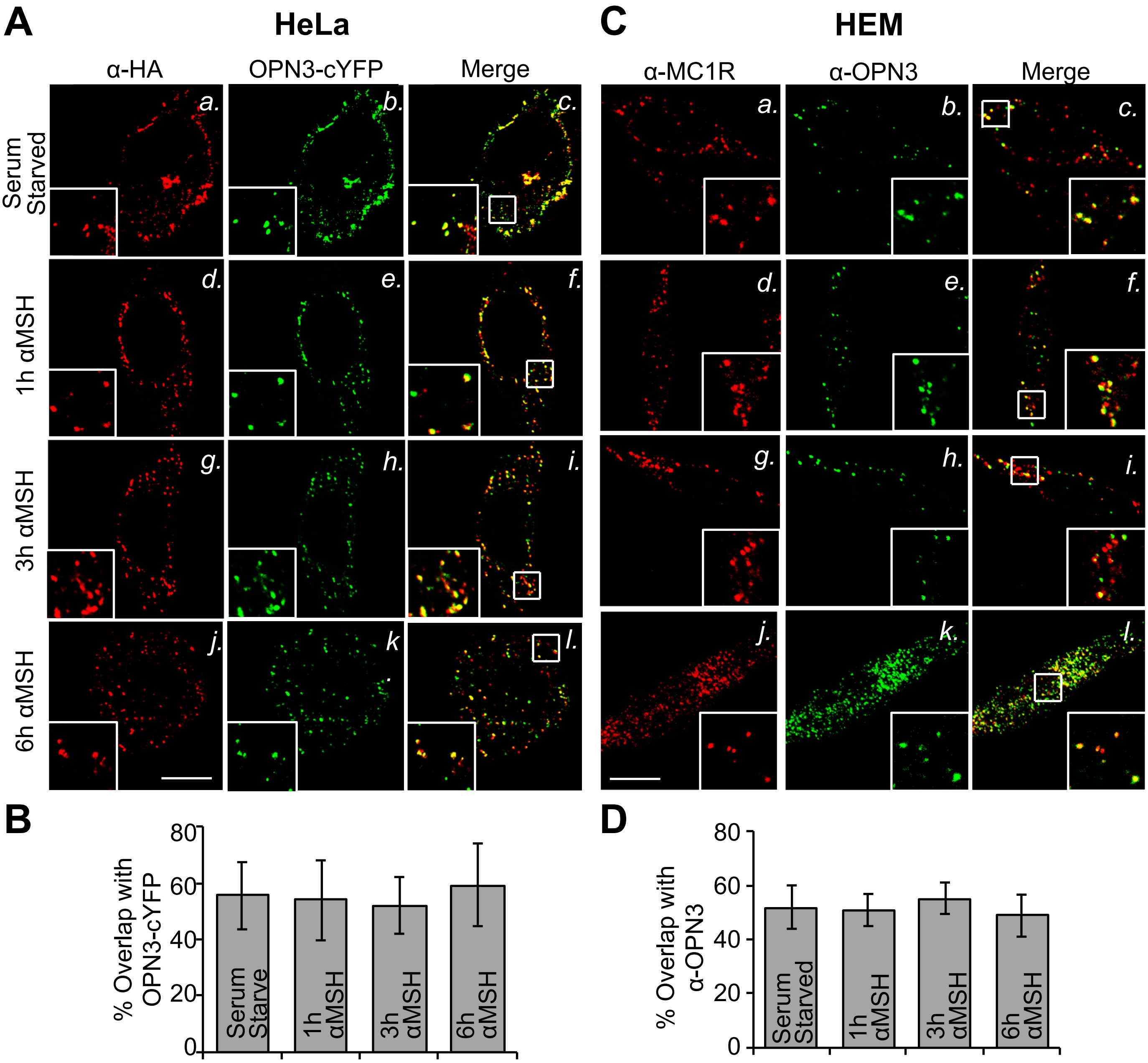


**Supp. Fig. 7. αMSH stimulation does not affect the colocalization of MC1R and OPN3.**

A. **Representative confocal images of HeLa cells coexpressing OPN3-cYFP and MC1R-nHA and immunostained with anti-HA antibody.** Cells were serum starved overnight and treated with vehicle (DMSO, < 0.1%), 0.5 μM αMSH for 1, 3, or 6 hours. Calibration bar, 10 μm.

B. **αMSH treatment does not affect colocalization of OPN3-cYFP and MC1R-nHA in Hela cells.** Bar graph representing the percent overlap between OPN3-cYFP and MC1R-nHA fluorescent signal in HeLa cells shows no significant difference between the absence and presence of αMSH for different time intervals. n=12 cells from 3 independent experiments; ± SEM.

C. **Representative confocal images of HEM cells immunostained with anti-MC1R and anti-OPN3 antibodies.** Cells were serum starved overnight or treated with 0.5 μM αMSH for 1, 3, or 6 hours. The final DMSO concentration did not exceed 0.1% (v/v). Calibration bar, 10 μm.

D. **Bar graphs representing the percent overlap between OPN3 and MC1R fluorescent signals of HEM cells.** n=9 cells from 3 independent experiments, ± SEM.

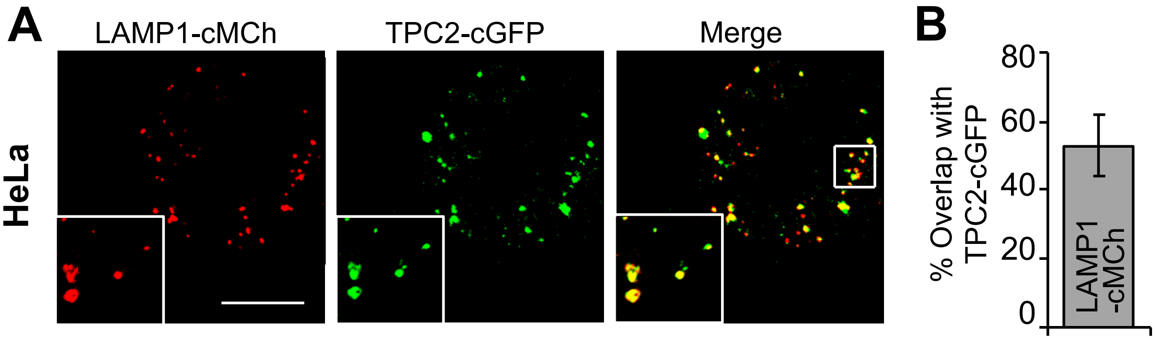


**Supp. Fig. 8. Colocalization analysis of the lysosomal proteins TPC2 and LAMP1 in HeLa cells.**

**A. Representative fluorescence confocal images of HeLa cells coexpressing TPC2-cGFP and LAMP1-cMCh.** Calibration bar, 10 µm.

**B. The percent overlap between LAMP1 and TPC2 fluorescent signals from A.** There is significant (~50%) overlap of LAMP1 and TPC2. n=15 cells from 3 independent experiments, ± SEM.
